# Trigeminal innervation and tactile responses in mouse tongue

**DOI:** 10.1101/2023.08.17.553449

**Authors:** Linghua Zhang, Maximilian Nagel, William P Olson, Alexander T. Chesler, Daniel H. O’Connor

## Abstract

The mammalian tongue is richly innervated with somatosensory, gustatory and motor fibers. These form the basis of many ethologically important functions such as eating, speaking and social grooming. Despite its high tactile acuity and sensitivity, the neural basis of tongue mechanosensation remains largely mysterious. Here we explored the organization of mechanosensory afferents in the tongue and found that each lingual papilla is innervated by Piezo2^+^ trigeminal neurons. Notably, each fungiform papilla contained highly specialized ring-like sensory neuron terminations that asymmetrically circumscribe the taste buds. Myelinated lingual afferents in the mouse lingual papillae did not form corpuscular sensory end organs but rather had only free nerve endings. *In vivo* single-unit recordings from the trigeminal ganglion revealed lingual low-threshold mechanoreceptors (LTMRs) with conduction velocities in the Aδ range or above and distinct adaptation properties ranging from intermediately adapting (IA) to rapidly adapting (RA). IA units were sensitive to both static indentation and stroking, while RA units had a preference for tangential forces applied by stroking. Lingual LTMRs were not directly responsive to rapid cooling or chemicals that can induce astringent or numbing sensations. Sparse labeling of lingual afferents in the tongue revealed distinct terminal morphologies and innervation patterns in fungiform and filiform papillae. Together, our results indicate that fungiform papillae are mechanosensory structures, while suggesting a simple model that links the functional and anatomical properties of tactile sensory neurons in the tongue.

## Introduction

The sense of touch is crucial for the function of the mammalian tongue in numerous behaviors including chewing (Pippi et al., 2018), swallowing (Chee et al., 2005; Tei et al., 2004), social grooming (Stern and Johnson, 1989) and vocalizing (Blom, 1960; Niemi et al., 2002). Unlike tactile information from the skin, tactile information from the tongue is often closely associated with chemo- and thermosensory inputs from food and drink, which together produce a vivid and complex mouthfeel. In terms of tactile acuity and sensitivity, the tip of the human tongue outperforms the fingertips with a lower two-point discrimination threshold and a lower mechanical threshold (Trulsson and Essick, 1997, 2010; Aktar et al., 2015; Miles et al., 2018). However, compared to other touch-sensitive organs including the hands/paws and the whiskers (Abraira and Ginty, 2013), the tongue has been less well characterized in terms of the sensory neuron subtypes and local architecture of nerve terminals underlying tactile sensitivity.

The dorsal surface of the tongue, which contacts food and drink, is covered with stratified squamous epithelium and contains four types of small protrusions called papillae: filiform, fungiform, circumvallate and foliate papillae, of which the filiform papillae and the fungiform papillae are the most abundant (Goździewska-Harłajczuk et al., 2018; Reginato et al., 2014) (**Figure S1**). On the mouse tongue, there are over 7000 conical-shaped filiform papillae that cover almost the entirety of the dorsal surface (Y. Wang et al., 2016). The less abundant, mushroom-shaped fungiform papillae host taste buds, and are concentrated at the tip and sides of the tongue. Based on their shape and abundance, filiform papillae have been hypothesized to be responsible for receiving tactile input (Böck, 1971; Kunze, 1969; Sato et al., 1988) and to act either as strain sensors or strain amplifiers (Lauga et al., 2016). However, the observation that sensory afferents expressing the mechanotransduction channel Piezo2 (Chesler et al., 2016; Ranade et al., 2014) can be found in both filiform and fungiform papillae (Moayedi et al., 2018), and that the density of fungiform papillae is positively correlated with lingual tactile sensitivity in humans (Bangcuyo and Simons, 2017; Essick et al., 2003; Yackinous and Guinard, 2001; Zhou et al., 2021), suggests that fungiform papillae may also be involved in touch sensation. Complicating the matter further, a group of fibers in the chorda tympani – a branch of the facial nerve that innervates taste buds within fungiform papillae and carries taste information – can also respond to stroking on the tongue (Donnelly et al., 2022; Donnelly et al., 2018). Questions about how the filiform and fungiform papillae are innervated by somatosensory neurons from the trigeminal ganglion (TG), how lingual LTMRs respond to different tactile stimuli, and whether those nerve terminals form sensory end organs in the tongue, remain to be answered.

Here, we approached these questions using single-unit recordings from TG lingual LTMRs in anesthetized mice together with genetic labeling and tracing experiments. Our results uncover two groups of lingual LTMRs that differ in their tactile response properties, and suggest a model in which these two groups of LTMRs innervate different types of papillae and convey different information about touch on the tongue.

## Results

### Trigeminal afferents innervate both filiform and fungiform papillae and have the highest density at the tip of the tongue

To investigate how papilla structures on the tongue are innervated by trigeminal neurons, we did anterograde tracing in fixed mouse tissue by placing DiI crystals into the TG and examined the entire dorsal surface of the tongue in mice from postnatal day (P)0 - P21 (n = 10). We found that both fungiform papillae (**Figure 1A** and **1B**, triangles; Figure **1C**) and filiform papillae (**Figure 1A** and **1B**, arrows; **Figure 1D**) were innervated by DiI-labeled trigeminal afferents, and that the tip of the tongue had the highest trigeminal innervation density across animals (**Figure 1E**). The efficiency of DiI tracing decreased at P21, as indicated by the lower signal-to-noise ratio in P21 in **Figure 1E**, with peaks of fluorescence representing the locations of fungiform papillae much higher in P21 compared to P10 and P12. Surprisingly, fungiform papillae, the taste-bud holding structures concentrated at the anterior region of the tongue, were significantly more heavily innervated by trigeminal afferents than the uniformly distributed filiform papillae (**Figure 1B**, cf. **Figure 2A**). In comparison, previous studies suggest that the chorda tympani afferents from the geniculate ganglion (GG), which consist of gustatory fibers primarily and a subset of thermo- or mechanosensory fibers (Yokota and Bradley, 2016; Donnelly et al., 2018), only innervate fungiform papillae (Fei et al., 2014). Together, these findings suggest that the innervation pattern of TG afferents differs from that of GG afferents, and that both types of lingual papillae may be involved in somatosensation (**Figure 1F**).

**Figure 1.**
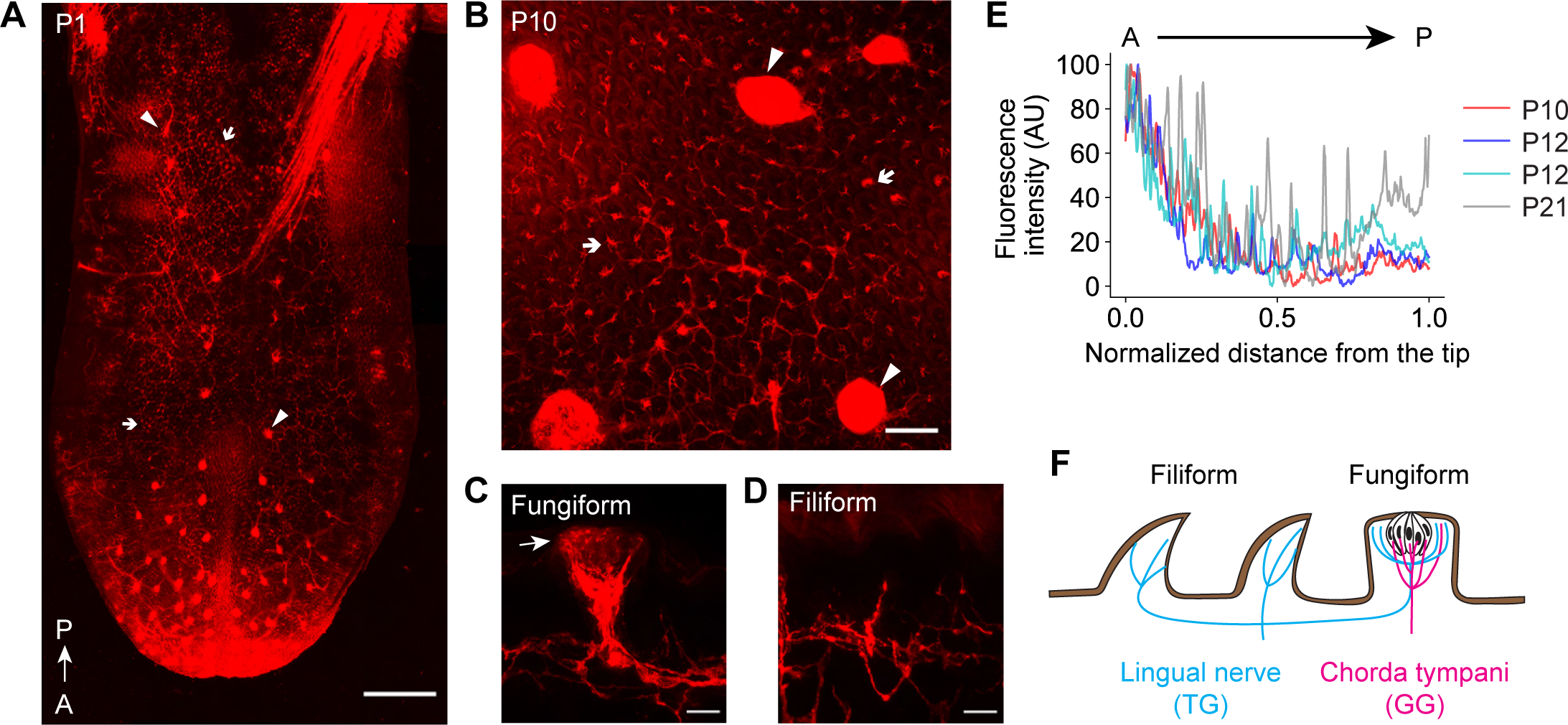
Trigeminal afferents innervate both filiform and fungiform papillae and have the highest density at the tip of the tongue. **(A-B)** Dorsal views of the tongue showing DiI-labeled trigeminal axons in P1 **(A)** and P10 **(B)** mice. Arrows: filiform papillae. Triangles: fungiform papillae. To reveal the fluorescent signals in the filiform papillae in **(B)**, those in the fungiform papillae were saturated due to contrast adjustments. Scale bar: 500 µm for **(A)** and 100 µm for **(B)**. **(C-D)** Longitudinal views of the tongue showing DiI-labeled trigeminal axons in fungiform **(C)** and filiform **(D)** papillae. The arrow in **(C)**: axons terminated at the extragemmal regions of a fungiform papilla. Scale bar: 20 µm. **(E)** The fluorescence intensity along the A-P axis across DiI-labeled tongue surfaces in different ages. The farthest normalized distance from the tip (1.0) is defined by the location of the last fungiform papilla at the posterior end. **(F)** A schematic of the innervation patterns of trigeminal ganglion (TG) neurons and geniculate ganglion (GG) neurons on filiform and fungiform papillae. Fungiform papillae are innervated by both chorda tympani nerve from GG and lingual nerve from TG, whereas filiform papillae are only innervated by lingual nerve from TG.

**Figure 2.**
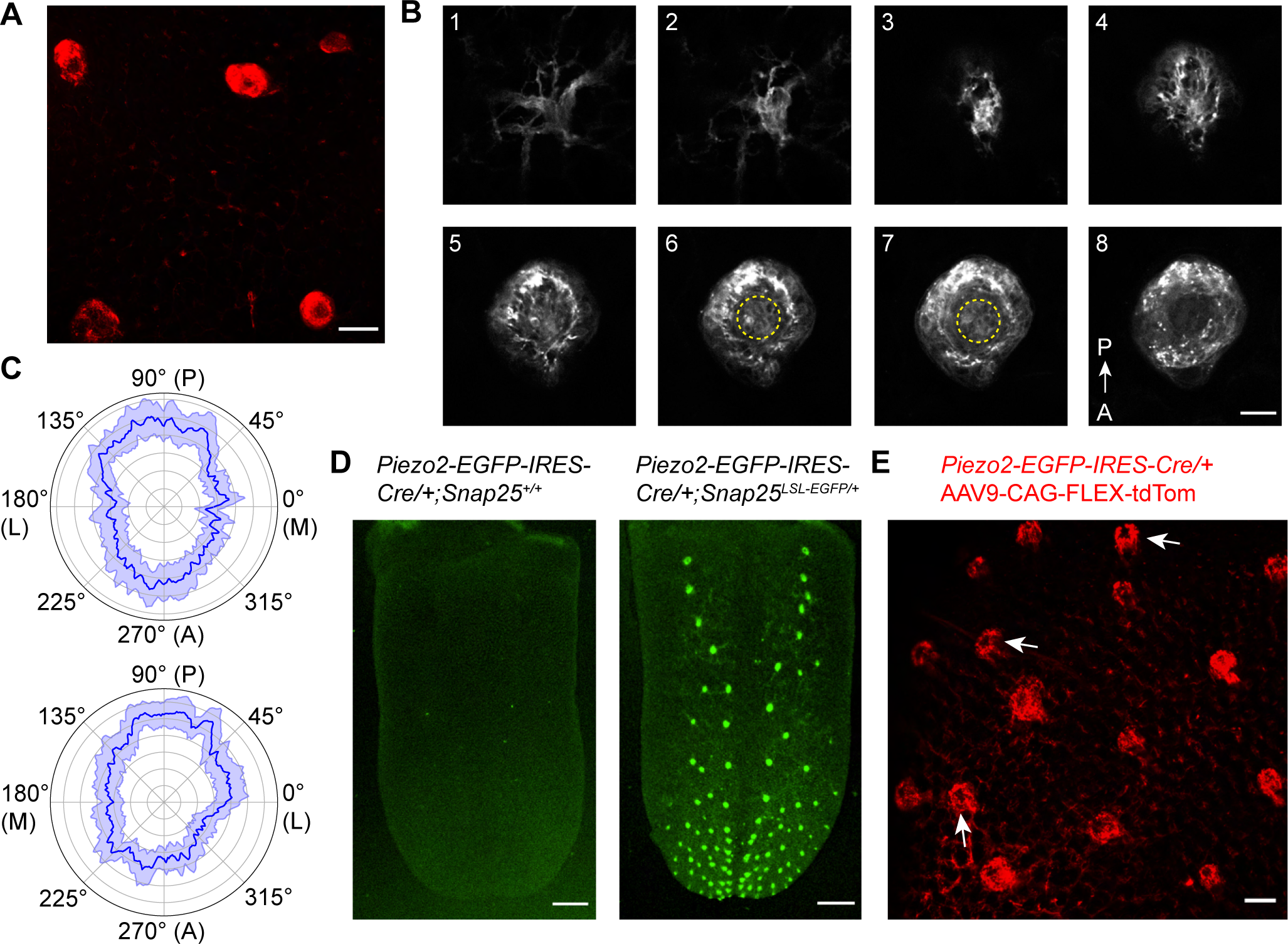
Trigeminal afferents asymmetrically innervate the extragemmal region of fungiform papillae and exhibit a ring-like termination pattern surrounding the taste pore, similar to Piezo2+ afferents. **(A)** An unsaturated maximum intensity projection of the confocal z-stack in Figure 1B showing the ring-like innervation pattern of DiI-labeled trigeminal afferents in fungiform papillae. Scale bar: 100 µm. **(B)** Optical sections of DiI-labeled trigeminal afferents in a fungiform papilla from a top-down view. The sections are labeled in an order from the deeper connective tissue (section 1) to the apical epithelium (section 8). Taste bud regions are circled in yellow dashed lines in sections 6 and 7. Scale bar: 25 µm. **(C)** The fluorescence intensity of DiI in the ring region of fungiform papillae at each side of the tongue (top and bottom) on a linear scale (gray concentric circles). The dark blue line represents the mean intensity across fungi-form papillae, while the light blue shading indicates the bootstrap 95% confidence interval. n = 26 and 27 papillae from 4 (2 P12 and 2 P21) mice for top and bottom, respectively. **(D)** Dorsal view of the tongue in *Piezo2-EGFP-IRES-Cre/+;Snap25^LSL-EGFP/+^* and its *Piezo2-EGFP-IRES-Cre/+; Snap25^+/+^* littermate. Scale bar: 500 µm. **(E)** In *Piezo2-EGFP-IRES-Cre/+* mice injected with AAV9-CAG-FLEX-tdTom, tdTomato-labeled Piezo2 expressing afferents were found to innervate both fungiform and filiform papillae. The Piezo2^+^ nerve terminals also innervated the extragemmal region of fungiform papillae and form a ring-like innervation pattern (arrows). Scale bar: 100 µm.

### Trigeminal afferents asymmetrically innervate the extragemmal region of fungiform papillae and exhibit a ring-like termination pattern surrounding the taste pore, similar to Piezo2^+^ afferents

The longitudinal view of a fungiform papilla in **Figure 1C** shows that trigeminal afferents in a fungiform papilla penetrated the apical epithelium surrounding the taste pore and terminated in the extragemmal (outside of taste buds) region. To further explore the terminal organization of trigeminal afferents in fungiform papillae, we performed DiI anterograde tracing from the TG and examined fungiform papillae in confocal z-stacks, in order to better appreciate how the axons traveled through the papillae. We found that the trigeminal fibers exhibit a ring-like termination pattern in fungiform papillae in maximal intensity projections of the confocal stacks (**Figure 2A**), with little DiI signal in the middle part of the papillae where taste pores and taste buds were located. The trigeminal nerve fibers innervating a fungiform papilla met at the base of the connective tissue and entered the connective tissue core through the middle as a bundle (**Figure 2B**, sections 1–3; **Video S1**). The clusters of nerve fibers then bifurcated into multiple branches, traveled towards the side walls of the papilla and avoided the taste bud (**Figure 2B**, sections 4–7), while still going upwards beyond the level of the taste bud until reaching the apical epithelium surrounding the taste pore, forming a ring-shaped terminal organization (**Figure 2B**, sections 8). This finding is consistent with the description of the trigeminal innervation pattern in fungiform papillae in a previous study (Suemune et al., 1992), and their termination at the superficial epithelium suggests that the fungiform papillae innervating trigeminal afferents are capable of detecting sensory stimuli, including mechanical stimuli, in an extremely sensitive manner. We also found that the distribution of trigeminal afferents was asymmetric along the fungiform circumference. In both left and right sides of the tongue, the innervation was more concentrated in the posterior region of the fungiform papillae than in the medial/lateral regions (**Figure 2C**, n = 4 mice, n = 26 and 27 papillae for each side of the tongue, respectively).

To investigate the innervation pattern of mechanosensitive afferents expressing Piezo2, we examined the dorsal surface of the tongue in *Piezo2-EGFP-IRES-Cre/+;Snap25^LSL-EGFP/+^*mice aged P1–P2 (n = 4) and *Piezo2-EGFP-IRES-Cre/+* mice injected with AAV9-pCAG-FLEX-tdTomato virus intraperitoneally at ages P1–P2 (n = 4) to avoid lineage problems caused by embryonic expression of Piezo2. As with DiI-labeled trigeminal afferents, fluorescently-labeled Piezo2^+^ afferents were found in both fungiform and filiform papillae but not their *Piezo2-EGFP-IRES-Cre/+;Snap25^+/+^* littermates (**Figure 2D**), suggesting that both types of papillae are involved in mechanosensation. Afferents expressing Piezo2 in fungiform papillae also exhibited a ring-like termination pattern, and innervated the extragemmal region rather than the intragemmal (inside taste buds) region (**Figure 2E**, arrows; cf. **Figure 3C**, bottom row, arrowhead), similar to DiI-labeled trigeminal afferents (**Figure 1C**). Considering that GG afferents are not the major contributor to extragemmal afferents in fungiform papillae (Donnelly et al., 2022; Ohman-Gault et al., 2017), our results suggest that TG afferents are the main mechanosensory supply for fungiform papillae.

**Figure 3.**
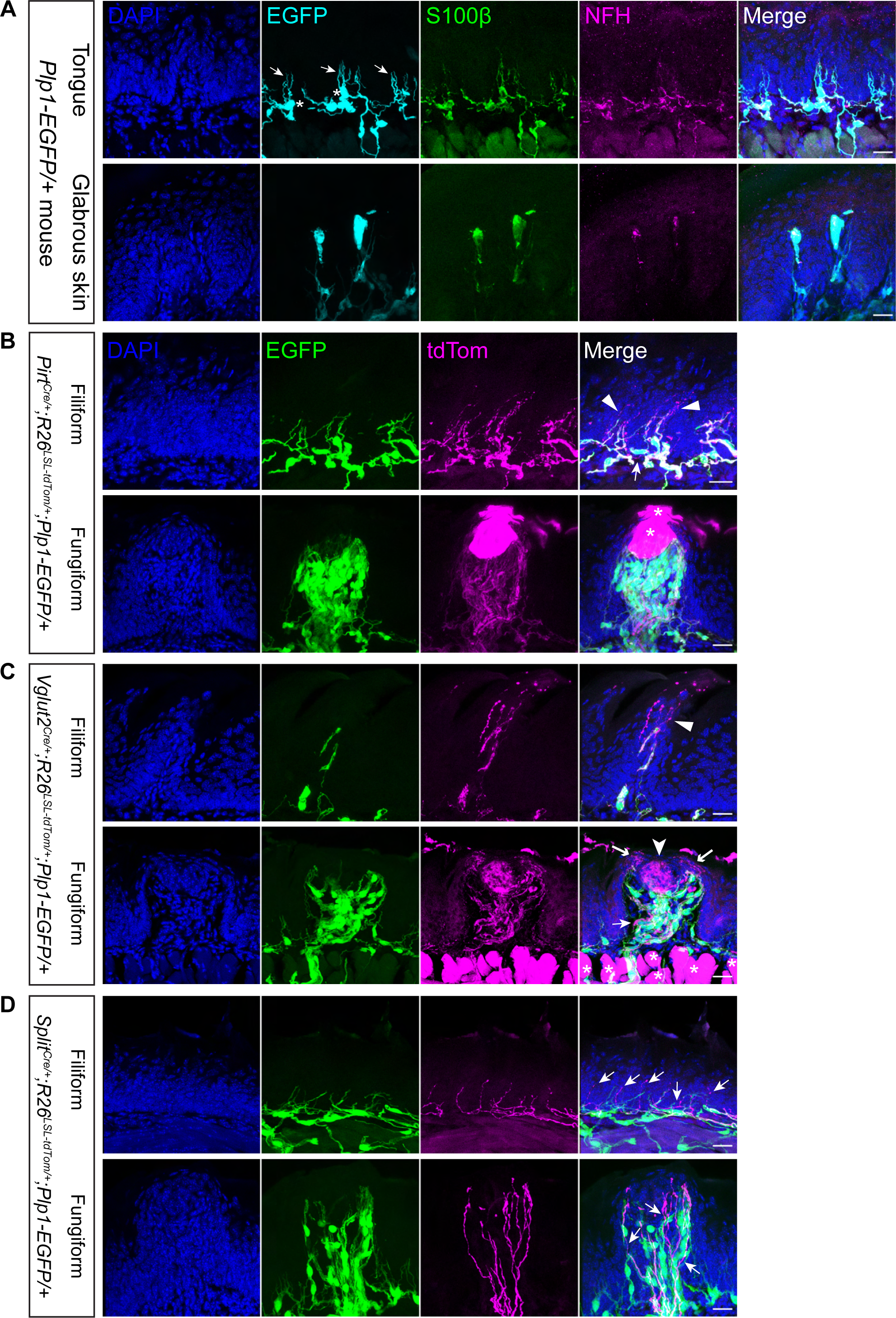
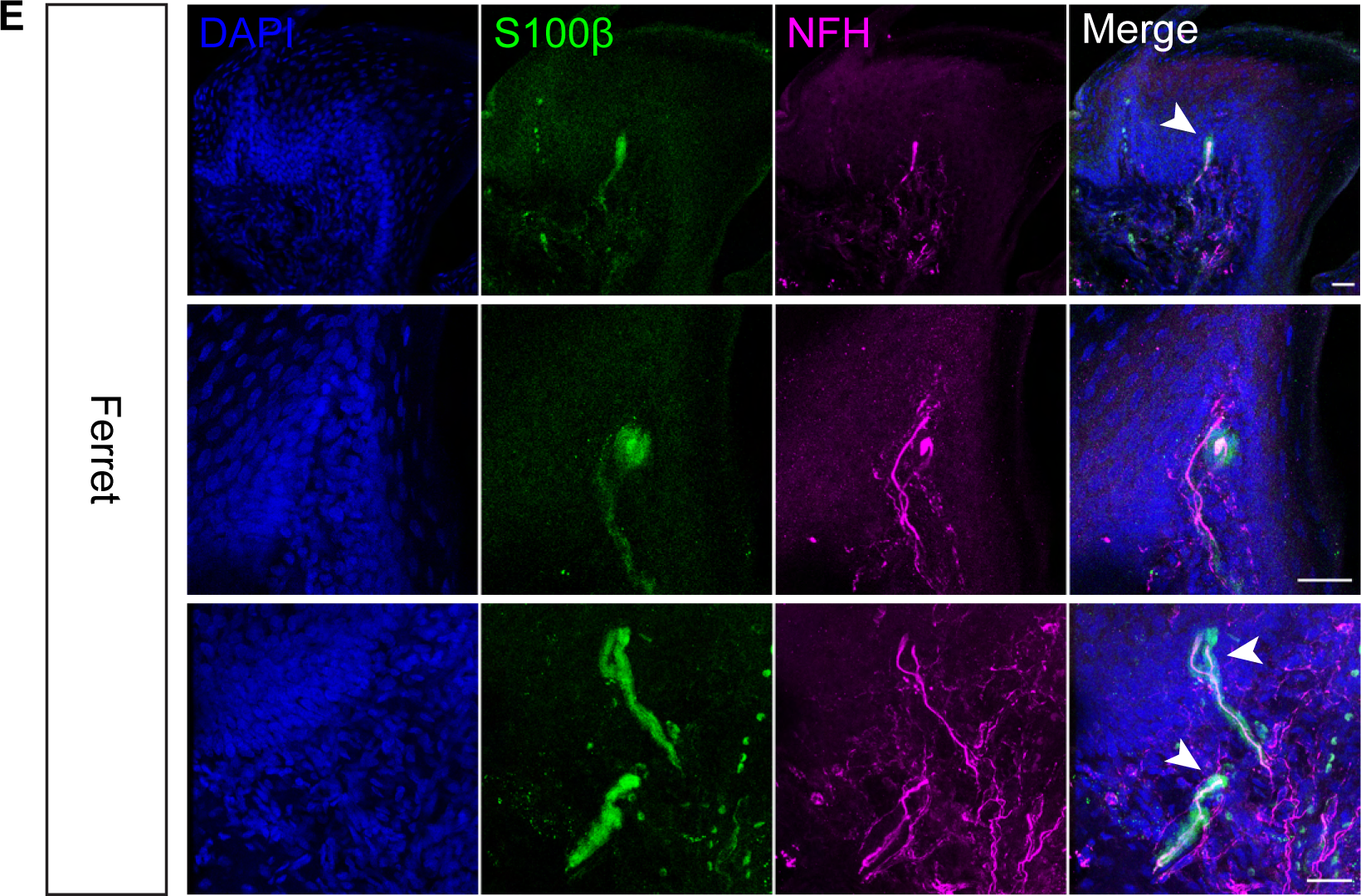
Mouse filiform papillae are innervated by NFH+ myelinated free nerve endings, while ferret filiform papillae contain corpuscular organs resembling end bulbs of Krause. **(A)** In the tongue of *Plp1-EGFP/+* mice, no corpuscular end organs but only myelinated free nerve endings can be identified in the connective tissue core of filiform papillae (top), compared to the glabrous skin, where Meissner corpuscles can be found in the dermal papillae (bottom). DAPI: cell nucleus. EGFP and S100β: (terminal) Schwann cells. NFH: myelinated nerve fibers. Asterisks: cell bodies of Schwann cells. Arrows: cell processes or myelin from Schwann cells. Scale bar: 20 µm. **(B)** In *Pirt^Cre/+^;R26^LSL-tdTom/+^;Plp1-EGFP/+* mice, tdTom-labeled, myelinated afferents that colocalized with EGFP can be seen in both filiform and fungiform papillae. Taste bud cells in and around fungiform papillae were also labeled by *Pirt^Cre^* (asterisks). The myelin was lost at the nerve terminal in myelinated afferents (triangles). *Pirt^Cre^* also labeled unmyelinated afferents (thin arrows). **(C)** In *Vglut2^Cre/+^;R26^LSL-tdTom/+^;Plp1-EGFP/+* mice, tdTom-labeled, myelinated afferents that colocalized with EGFP can be seen in both filiform and fungiform papillae. Muscles were also labeled by *Vglut2^Cre^* (asterisks). In fungiform papillae, *Vglut2^Cre^* labeled intragemmal (arrowhead) and extragemmal (thick arrows) afferents, both of which were not myelinated at the nerve terminals. *Vglut2^Cre^* also labeled unmyelinated afferents (thin arrows). **(D)** *Split^Cre^* labeled unmyelinated afferents in both filiform and fungiform papillae (arrows). Scale bars in **(B-D)**: 20 µm. **(E)** In ferret tongue, corpuscular end organs with cylindrical (top, bottom) or globular (middle) shapes resembling end bulbs of Krause can be seen in the connective tissue core or the basal layer of epithelium in filiform papillae, usually with blunt, enlarged nerve endings (arrowheads). The structure of filiform papillae is more complex in ferrets, as there can be more than one connective tissue papillae inside a single filiform papilla (top). Scale bar: 25 µm.

### Mouse filiform papillae contain NFH^+^ myelinated free nerve endings without sensory end organs, while ferret filiform papillae contain corpuscular end organs resembling end bulbs of Krause

Sensory end organs are specialized structures formed at the terminals of sensory afferents that can act as mechanical filters and thus affect the mechanosensory responses of the axons. We next sought to describe any sensory end organs in the mouse tongue, with a special focus on the filiform papillae, where sensory innervation is solely provided by trigeminal afferents.

We found that both filiform papillae and fungiform papillae in the mouse tongue are innervated by NFH^+^ myelinated afferents, consistent with prior work (Moayedi et al., 2018; Sato et al., 1988) (**Figure 3A** and **S2**). In order to visualize the shape of terminal Schwann cells, which are often involved in the structure of sensory end organs, we used both immunostaining (S100β) and genetic labeling (*Plp1-EGFP*) methods targeting terminal Schwann cells. In contrast to the glabrous skin, where we easily identified corpuscular end organs such as Meissner corpuscles by immunostaining S100β or genetically labeling PLP1 (**Figure 3A**, lower panels), we could not find corpuscular end organs (e.g. structures resembling end bulbs of Krause or Meissner corpuscles in shape) in the filiform papillae of the mouse tongue (**Figure 3A**, upper panels). The lack of Merkel cells at the apical epithelium of tongue reported by other research groups (Donnelly et al., 2022; Moayedi et al., 2018) also suggests that the nerve terminals ascending the connective tissue cores of papillae were not likely to be associated with Merkel cells.

By conducting a small-scale screen of Cre mouse lines (**Table 1**), we identified several Cre lines that can label neuronal afferents on the surface of the tongue, including *Pirt^Cre^*, *Vglut2^Cre^* and *Split^Cre^*, which allowed us to better visualize the afferent terminals with genetic labeling strategies. In both *Pirt^Cre^;Plp1-EGFP;R26^LSL-tdTomato^*mice and *Vglut2^Cre^;Plp1-EGFP;R26^LSL-tdTomato^* mice, tdTomato-labeled, myelinated as well as unmyelinated primary afferents can be found in both fungiform papillae and filiform papillae. The terminal Schwann cells in mouse filiform papillae did not arrange in a compact stack and form either oval or cylindrical structures overall as in sensory corpuscles; rather, they merely formed a myelin sheath, which went missing before reaching the terminations of lingual afferents, leaving the nerve endings free (**Figure 3B** and **3C**, triangles). In fungiform papillae, *Vglut2^Cre^* labeled at least two different groups of afferents defined by their terminal locations: the intragemmal afferents innervating the taste bud (**Figure 3C**, arrowhead), and the extragemmal afferents that penetrated the apical epithelium surrounding the taste pore (**Figure 3C**, thick arrows). The terminals of both fungiform groups also did not associate with any PLP1-EGFP^+^ corpuscular structures. In *Split^Cre^;Plp1-EGFP;R26^LSL-tdTomato^* mice, tdTomato-labeled unmyelinated primary afferents with free nerve endings were found in both fungiform papillae and filiform papillae (**Figure 3D**, arrows), which may represent nociceptors with lower conduction velocities.

**Table 1.** Genetic screening of Cre driver lines that can label lingual afferents (crossed with R26*^LSL-tdTom^*)

| Genotype | Reported labeling of sensory neuronal subtypes | References | Tamoxifen amount and application time in this study | Expression pattern in the anterior tongue identified in this study |
| --- | --- | --- | --- | --- |
| *TrkB<sup>CreERT2</sup>* | Aδ LTMRs in DRG and lamellar cells in human Meissner corpuscles | Calavia et al., 2010; Li et al., 2011; Rutlin et al., 2014 | 0.1 mg @ P0 | Flat-shaped cells in the epithelium (could be areas adjacent to fungiform papillae) and cells with dendritic protrusions in the lamina propria |
| *TrkC<sup>CreERT2</sup>* | Proprioceptors and Merkel afferents | Bai et al., 2015; Severson et al., 2017 | 3 mg @ E12.5; 0.1 mg @ P5 and P6, respectively | Cells in the connective tissue underneath filiform papillae; in E12.5, NFH-negative fibrous structures in deeper muscles were labeled, and some of them were also PGP9.5-negative |
| *Etv1<sup>CreERT2</sup>* | Proprioceptors; Pacinian corpuscle-innervating DRG neurons; Non-myelinating Schwann cells of Pacinian and Meissner corpuscles | de Nooij et al., 2013; Fleming et al., 2016; Prsa et al., 2019 | 0.1 mg @ P6 - P10 | Cells in filiform and fungiform papillae |
| *Vglut1<sup>IRES-cre</sup>* | Proprioceptors and Aβ SA-LTMRs | Brumovsky et al., 2007 | N/A | Muscles in the tongue; Nerve fibers (very sparse) |
| *Advillin<sup>Cre</sup>* | Primary sensory neurons in DRG and TG | Lopes et al., 2017; Zurborg et al., 2011 | N/A | Flat-shaped cells in the epithelium of filiform papillae (different from those in *TrkB<sup>CreERT2</sup>*); Cells with dendritic protrusions in the lamina propria |
| *PV<sup>Cre</sup>* | Proprioceptors; Pacinian, Meissner and | de Nooij et al., 2013; Wu | N/A | Muscles in the tongue (different from |
|  | lanceolate endings of DRG neurons; TG neurons innervating the tongue | et al., 2018 |  | *Vglut1*<sup>IRES-cre</sup> patterns); Occasionally labels thick fibrous structures that very likely consisted of rod-shaped cells in the lamina propria; Nerve fibers, including extragemmal afferents in fungiform papillae. |
| *TH*<sup>Cre</sup> | C-LTMRs and sympathetic nerves | Li et al., 2011 | N/A | Hard to identify individual nerves after 3 days' 6-OHDA treatment |
| *MafA*<sup>Cre</sup> | LTMRs in DRG | Bourane et al., 2009 | N/A | Muscles in the tongue |
| *Pirt*<sup>Cre</sup> | Sensory neurons (including GG neurons) | Kim et al., 2008 | N/A | Nerve fibers in both fungiform and filiform papillae; Cells within and near the taste buds |
| *Vglut2*<sup>Cre</sup> | GG neurons; nociceptors in DRG | Lagerström et al., 2010; Vandenbeuch et al., 2010 | N/A | Nerve fibers in both fungiform and filiform papillae |
| *Split*<sup>Cre</sup> | Aβ RA-LTMRs in hairy skin and whisker pad (DRG and TG) | Rutlin et al., 2014; unpublished data | N/A | Unmyelinated afferents in filiform and fungiform papillae |

The average size and the shape of tongue filiform papillae vary across mammalian species, as do their degrees of keratinization and their structural complexity. Previous studies have identified the existence of a type of simple corpuscle through immunohistochemistry and electron microscopy in the more developed filiform papillae of cats and cattle (Sato et al., 1988; Spassova, 1974), but not in the smaller and simpler filiform papillae of rats (Sato et al., 1988). Contrary to previous findings suggesting the existence of a type of simple corpuscle called Krause end bulbs in mouse tongue (Moayedi et al., 2018), we could only find unencapsulated nerve endings in both types of papillae in mice. To provide comparative context and test our ability to identify lingual end organs, we examined nervous elements in the ferret tongue by immunostaining myelinated afferents (NFH) and terminal Schwann cells (S100β). We found that the majority of nerve afferents in the ferret tongue were not associated with corpuscles. However, unlike in the mouse tongue, ferret filiform papillae contained end organ structures with varied shapes, including cylindrical or globular structures that resembled the simple end bulbs of Krause in cats (**Figure 3E**, top and middle panels). Individual filiform papillae could contain more than one end organ (**Figure 3E**, bottom panel). Moreover, the lingual afferents encapsulated in a corpuscle usually had enlarged, blunt nerve endings (**Figure 3E**, all three panels), and could branch at the terminal (**Figure 3E**, bottom panel). Collectively, these results demonstrate that our methods allow us to identify sensory end organs, highlight structural differences among filiform papillae of different animal species, and show that those of the mouse lack obvious sensory end organs.

### Single-unit recordings reveal lingual LTMRs ranging from rapidly to intermediately adapting with varied tactile response properties and conduction velocities in the A-fiber range

We first determined the location of tongue-innervating neurons in TG by retrograde DiI tracing from the tip of the tongue. We found that TG neurons innervating the tip of the tongue were mostly located near or at the ventral side of the TG mandibular branch rather than the dorsal side (**Figure S3A**), as significantly lower numbers of tongue neurons (soma size of ≥20 µm) could be visualized at the dorsal TG by a one-photon fluorescence microscope (**Figure S3B**, dorsal: 14.9 ± 14.2, ventral: 71.3 ± 40.4; p = 0.0011, paired t-test; n = 9 TG from 5 mice). The ventral location of the tongue neurons within the TG makes them amenable to single-unit recordings but possibly challenging targets for optical monitoring. In contrast, whisker pad-innervating TG neurons could be found in abundant numbers at both the dorsal and ventral sides of TG (**Figure S3A**).

To make single-unit recordings, the tongue of an anesthetized mouse was carefully pulled out of the mouth using blunt forceps and placed on a surface that had a stable slope of 30 degrees from the vertical and against which the lower teeth were braced to keep the mouth open during recordings (**Figure 4A**). The surface of the tongue was gently stimulated using a fine-tip brush until the activity of a unit that was responsive to brush strokes was captured by the electrode in the TG. The stable platform setup prevented tongue stimulation from causing unintended, correlated movements of the jaw and the lower chin as well as any potential mechanical stimulation of facial hairs on the lower face, which could lead to spurious inclusion of non-tongue units.

**Figure 4.**
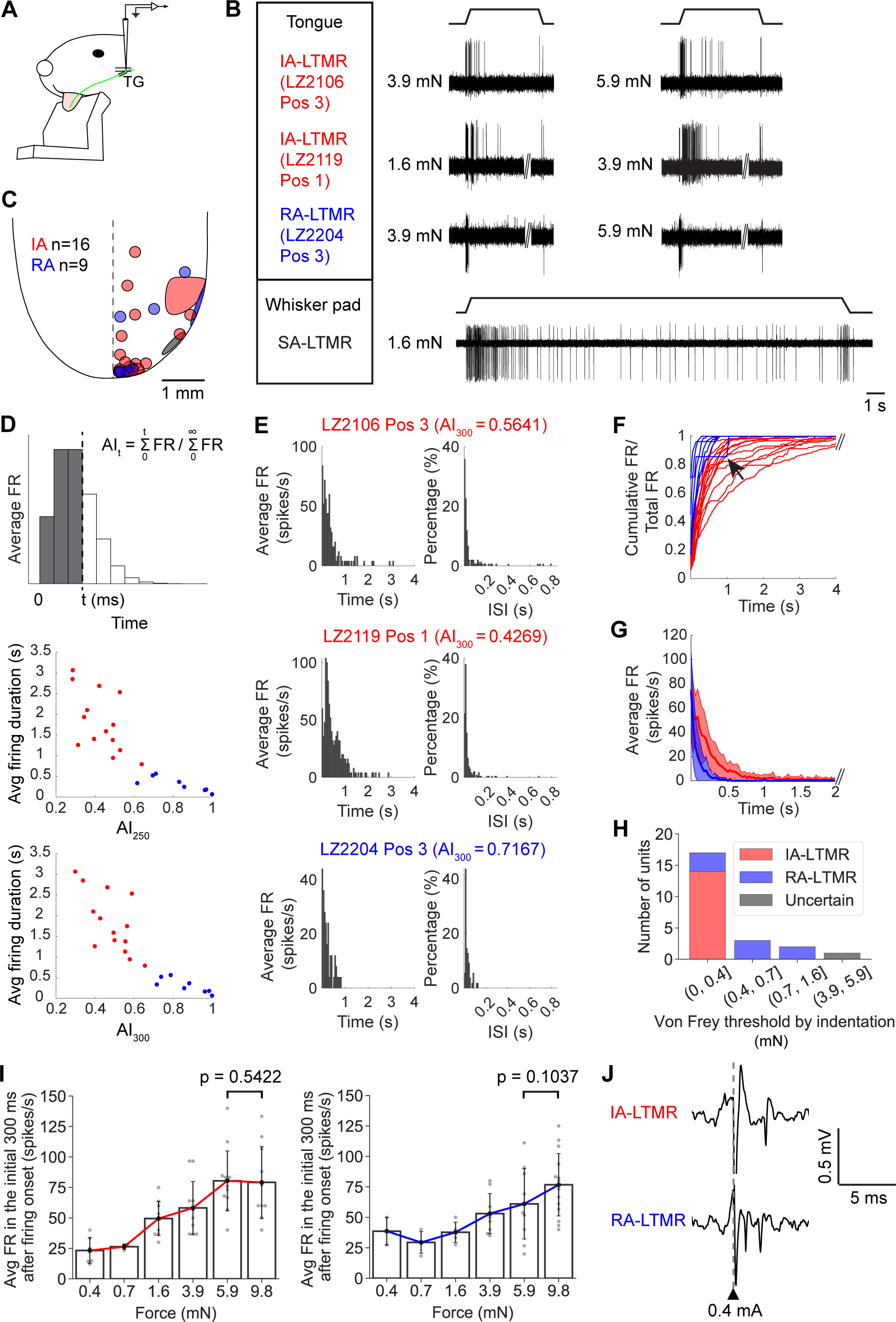
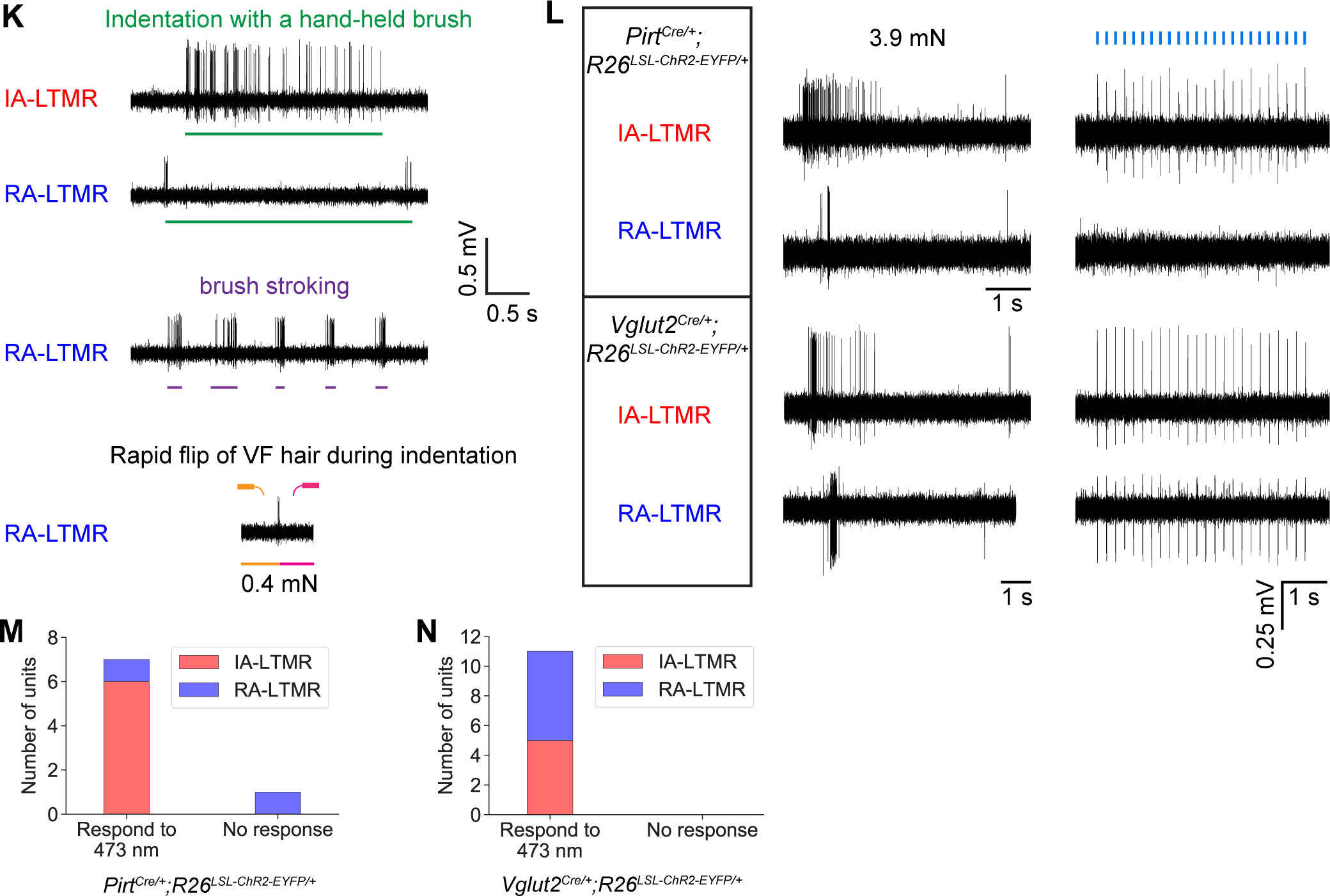
Single-unit recordings reveal lingual LTMRs ranging from rapidly to intermediately adapting with varied tactile response properties and conduction velocities in the A-fiber range. **(A)** Schematic of the single-unit recording setup. Green: lingual nerve. **(B)** Neural activities of single lingual LTMR units recorded in TG in response to tactile indentation exerted at their receptive fields at the tongue surface. Neural traces with no spikes were cropped in order to make the onsets and offsets of indentation aligned for each column. An example SA-LTMR innervating the whisker pad was included as a comparison. **(C)** The receptive fields of lingual LTMRs mapped by von Frey hairs (n = 25 from 19 mice; 16 IA-LTMRs marked in red, 9 RA-LTMRs in blue and 1 unit that cannot be categorized into IA or RA in gray). **(D)** The adaptation index (AI) and average firing duration of IA/RA-LTMRs in 3.9 mN stimulation. Top: schematic showing the calculation of AI_t_ using a peristimulus time histogram (PSTH) generated from a Poisson distribution (λ = 2). **(E)** PSTHs (bin width: 50 ms, n = 5 events) and interspike intervals (ISIs, bin width: 10 ms) of the 3 example lingual units in **B** in response to 3.9 mN. **(F)** The ratio of cumulative firing rate over total firing rate in IA/RA-LTMRs during 3.9 mN stimulation. Traces later than 4s were cropped. Arrow: a RA-LTMR with extremely low firing rates (LZ2203 Pos 6 Unit 3 in **Figure S4B**). **(G)** The average firing rate of IA/RA-LTMRs during 3.9 mN stimulation (trace and shading: mean ± SD). **(H)** The von Frey indentation threshold of lingual LTMRs (n = 23 from 16 mice; 14 IA-LTMRs, 8 RA-LTMRs and 1 unit that cannot be categorized into IA or RA). **(I)** The average firing rates of an example IA-LTMR (left) and a RA-LTMR (right) in the initial 300 ms after firing onset during von Frey indentation (mean ± SD. One-sided Student’s t-test). **(J)** Example neural traces in conduction velocity tests. The gray dashed line indicates the timepoint when a 0.4 mA electrical pulse was given to the unit’s receptive field. **(K)** The activities of lingual LTMRs in response to other tactile stimuli. The bottom row shows a RA-LTMR with a von Frey threshold higher than 0.4 mN. **(L)** Neural activities of example single lingual LTMR units recorded in TG in response to von Frey indentation and 473 nm light stimulation at their receptive fields in *Pirt^Cre/+^;R26^LSL-ChR2-EYFP/+^* and *Vglut2^Cre/+^;R26^LSL-ChR2-EYFP/+^* mice. Blue bars: 10 ms-duration 473 nm light pulses shone at 5 Hz. **(M-N)** Number of lingual LTMR units that can or cannot be activated by 473 nm light stimulation from 3 *Pirt^Cre/+^; R26^LSL-^ _ChR2-EYFP/+_* _mice and 8 *Vglut2*_*_Cre/+;R26LSL-ChR2-EYFP/+_* _mice._

Using von Frey filaments to apply a constant force on their receptive fields (RFs), we found lingual LTMRs with different adaptation properties, ranging from the rapidly adapting (RA) to intermediately adapting (IA). Except for a few units that exhibited low spontaneous activity, none of the tongue units we recorded behaved like classic slowly adapting (SA) units that could fire continuously from touch onset to offset, such as those in the whisker system (**Figure 4B**). As all tongue units showed a certain amount of adaptation to touch stimuli, here we define RA-LTMRs in the tongue to be those that stopped firing roughly within 0.5 s in response to all tested von Frey stimuli (*Methods*), and IA-LTMRs otherwise. We found the majority of lingual LTMRs had small RFs (less than 0.1 mm^2^). Most LTMRs recorded had their RFs located either within 0.5 mm of the tip of the tongue (14 out of 26, or 53.8%, **Figure 4C**) or along the midline or edge of the tongue, resembling the distribution of taste buds (cf. **Figure S1A** and **1A**).

We examined the adaptation properties of the tongue LTMR population by comparing their average firing duration and adaptation index (AI) under a static force of 3.9 mN. Here, the AI was defined as the cumulative firing rate within an initial time period from touch onset divided by the total cumulative firing rate. The closer the AI is to 1, the faster the unit adapts, indicating a more rapid decrease in the firing rate over time. We observed that the level of adaptation across the neuronal population could be seen as a continuum (**Figure 4D-F** and **S4A-B**), and that the IA and RA units had similar firing rates at the stimulus onset but different speeds of adaptation in later phases (**Figure 4G**).

Apart from the adaptation speed, lingual LTMRs also differed in their tactile sensitivity. Although lingual LTMRs in general had very low thresholds for static indentation, often less than 1.6 mN, RA-LTMRs usually had higher von Frey thresholds than IA-LTMRs. The thresholds for IA-LTMRs were consistently below 0.4 mN (**Figure 4H**), with some units registering as low as 0.2 mN (5 units) or 0.08 mN (1 unit, data not shown). Lingual LTMRs generally exhibited an elevated initial firing rate with increasing static force application. However, the IA-LTMRs, which were more sensitive, tended to reach saturation in their firing rates at lower forces compared to the less sensitive RA-LTMRs (**Figure 4I**).

To measure the conduction velocity of lingual LTMRs, we first estimated the distance between the anterior part of the tongue to the TG through lingual nerve dissection and retrograde DiI tracing. By exposing the lingual nerve posterior to its exit from the tongue and cutting the nerve anterior to its entrance at the foramen ovale, we estimated an upper bound of the distance from the anterior part of the tongue to the TG in adult mice to be 1.8 cm. By injecting DiI at the tip of the tongue in live mice and using the known DiI traveling speed *in vivo* (6 mm/day; Godement et al., 1987; Honig and Hume, 1986), we estimated a lower bound of the distance from the anterior part of the tongue to the TG in adult mice to be 1.1 cm. These distances are short enough to limit proper measurement of fast axonal conduction velocity due to potential masking by electrical artifacts of the first induced neuronal spike during stimulation. Hence, we could only estimate the conduction velocity of lingual LTMRs as falling into the range of Aδ fibers (lower bound: 6.1 ± 2.7 m/s, higher bound: 9.9 ± 4.5 m/s; n = 4. **Figure 4J**) or above.

Lingual LTMRs were generally highly responsive to brush stroking, although their sensitivity to vertical force changes varied across the population. IA-LTMRs were highly sensitive to vertical forces as indicated by their extremely low von Frey thresholds. They were also sensitive to subtle force variations in the vertical direction. When indented by a hand-held brush, where the applied force was not constant, unlike with von Frey filaments, they could maintain firing throughout the entire stimulation period, resembling SA-LTMRs. In contrast, RA-LTMRs fired only at the onset and offset of the indentation when stimulated with a hand-held brush (**Figure 4K**). RA-LTMRs were also sensitive to rapid changes of the buckling direction of von Frey filaments during indentation, even when the von Frey filaments used were below their von Frey thresholds for static indentation (**Figure 4K**). These observations suggest that RA-LTMRs were more sensitive to shear/tangential forces than to vertical forces.

As Pirt and Vglut2 were reported markers for geniculate ganglion neurons and primary sensory neurons including non-LTMR cell types in both trigeminal and dorsal root ganglion (Kim et al., 2008, 2016, 2015; Lagerström et al., 2010; Leijon et al., 2019; Patel et al., 2011; Vandenbeuch et al., 2010; Wu et al., 2015), we tested whether the neuronal afferents labeled by *Pirt^Cre^* or *Vglut2^Cre^* on the surface of the tongue partly originated from TG and included LTMR afferents. We performed single-unit recordings in the TG of *Pirt^Cre^;R26^LSL-ChR2^* mice and *Vglut2^Cre^;R26^LSL-ChR2^* mice, and recorded the responses of lingual LTMRs when their receptive fields were stimulated by 473 nm blue light (**Figure 4L**). 87.5% of lingual LTMRs in *Pirt^Cre^;R26^LSL-ChR2^* mice (6 IA-LTMRs and 1 RA-LTMR from 3 mice; **Figure 4M**) and 100% of lingual LTMRs in *Vglut2^Cre^;R26^LSL-ChR2^*mice (5 IA-LTMRs and 6 RA-LTMRs from 7 mice; **Figure 4N**) could be activated by blue light, suggesting that besides labeling GG neurons, *Pirt^Cre^* and *Vglut2^Cre^* can label lingual LTMRs in TG as well. Together, our results report that *Pirt^Cre^* and *Vglut2^Cre^* can label myelinated lingual afferents whose cell bodies are located in the TG, and that *Vglut2^Cre^*can label for both lingual IA- and RA-LTMRs.

### Lingual neurons that terminate in different types of papillae exhibit distinct terminal shapes and branching patterns

To investigate the terminal field patterns of lingual neurons in TG, we injected either AAVPHP.S- CAG-FLEX-tdTom or rAAV1-CAG-FLEX-tdTom virus into the TG in *Vglut2^Cre^* mice (n = 18), which allowed us to label a subset of mandibular branch neurons, including lingual LTMRs, in a series of labeling densities from sparse to dense (**Figure 5A**). In densely labeled samples, we found that the fungiform papillae from the anterior to the posterior part of the tongue were more heavily labeled by tdTomato compared to filiform papillae, and that the tip of the tongue, where fungiform papillae had the highest density, was thus equipped with the most TG afferents (**Figure 5B**). By sparsely labeling only a few lingual neurons, we identified at least 3 types of terminals with various shapes that innervated different parts of the tongue. At the dorsal side of the tongue, TG afferents innervating filiform papillae showed distinct branching patterns compared to those innervating fungiform papillae. Specifically, a single TG afferent projected to a single fungiform papilla and branched inside the papilla, whereas those innervating filiform papillae branched into multiple filiform papillae located nearby. At the junction of dorsal and ventral epithelium where no papilla structures were present, we observed small circular endings with no branches (**Figure 5C**). We did not attempt to trace the terminals of small-diameter afferents that might represent unmyelinated nociceptors.

**Figure 5.**
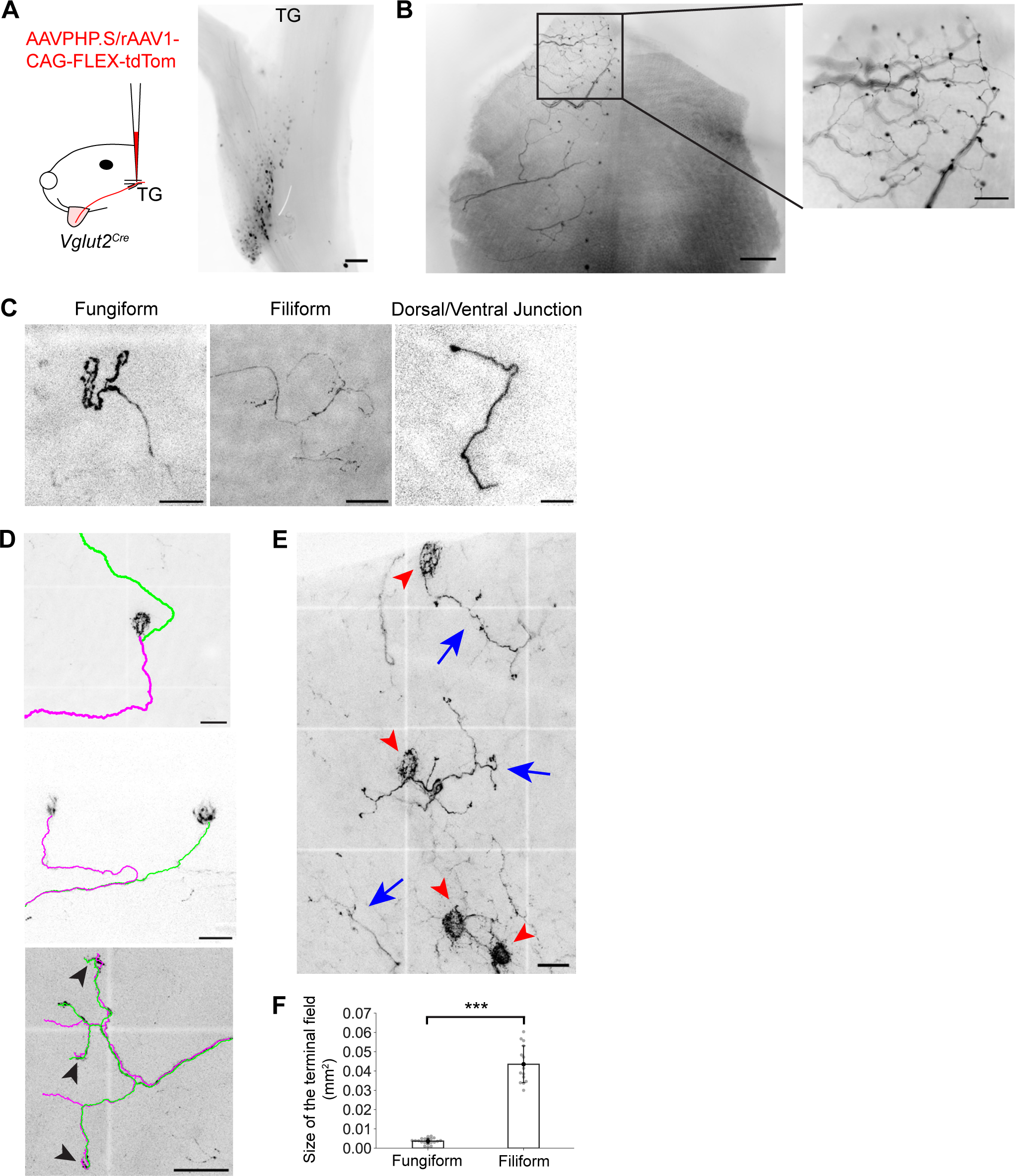
Lingual neurons that terminate in different types of papillae exhibit distinct terminal shapes and branching patterns. **(A)** Left: schematic of unilateral viral injection into the mandibular branch of the left TG in *Vglut2^Cre^* mice. Right: neurons at the mandibular branch of the TG were labeled by tdTom. Scale bar: 300 µm. **(B)** tdTom-labeled trigeminal afferents at the tongue surface. Only the left side of the tongue was labeled by unilateral viral injection. Scale bar: 800 µm. Inset: A zoomed-in view of the tip of the tongue showing many fungiform papillae strongly labeled by tdTom. Scale bar: 300 µm. **(C)** Single TG afferent terminals sparsely labeled by tdTom. Scale bar (left to right): 50 µm, 100 µm and 50 µm. **(D)** Multiple TG afferents simultaneously labeled by tdTom. Top: two clusters of nerve fibers (green & magenta) co-innervating a single fungiform papilla. Scale bar: 100 µm. Middle: two independent afferents (green & magenta) innervating two separate fungiform papillae. Scale bar: 50 µm. Bottom: two independent afferents (green & magenta) innervating several filiform papillae had overlap in their terminal fields. Arrowheads: their endings in the same filiform papillae. Scale bar: 100 µm. **(E)** Clusters of filiform papillae innervated by separate TG afferent aggregates (blue arrows). Red arrowheads: fungiform papillae. Scale bar: 100 µm. **(F)** The terminal field size of TG afferents innervating fungiform (0.0036 ± 0.0015, n = 20 from 5 mice) or filiform papillae (0.0435 ± 0.0096, n = 14 from 6 mice). p = 9.59 × 10^-10^, two-sided Welch’s t-test.

We further looked at the relationship of TG afferents innervating the same or nearby papillae structures in samples where more than a few afferents were labeled. We observed that multiple afferents innervating the same fungiform papillae could traverse the tongue in separate clusters of nerve fibers before reaching the papillae, and that afferents within the same nerve bundle could innervate different fungiform papillae (**Figure 5D**, top and middle). Notably, we did not find afferents that projected to multiple fungiform papillae by branching early before reaching them. In comparison, afferents innervating filiform papillae could travel in the same nerve bundle and exhibit overlap in their projected targets, while having different branching patterns (**Figure 5D**, bottom). The filiform papillae co-innervated by one or a few afferents formed distinct clusters (**Figure 5E**). The sizes of such filiform clusters were significantly larger than the terminal fields of fungiform-innervating afferents (fungiform: 0.0035 ± 0.0015 mm^2^, n = 20 from 5 mice; filiform: 0.0435 ± 0.0096 mm^2^, n = 14 from 6 mice; p = 9.59 × 10^-10^, two-sided Welch’s t-test; **Figure 5F**), suggesting a more restricted receptive field for the latter.

### Lingual LTMRs are insensitive to rapid cooling or chemicals that can induce astringent or numbing sensations

The mammalian tongue is a sensory surface where mechanosensation, thermosensation and chemosensation closely interact with each other. Our daily experience of eating and drinking reminds us that the perception of food texture can be affected by the food temperature (Engelen et al., 2003). Also, there are some interesting forms of tactile perception that can be induced by certain chemicals in food, such as the sense of astringency caused by chemicals containing the galloyl groups, including tannic acid in green tea, red wine and unripe fruits (Schöbel et al., 2014), and the sense of numbing induced by hydroxy-alpha-sanshool in green Sichuan peppercorns harvested from the *Zanthoxylum* species (Luo et al., 2022). We tested whether lingual LTMRs can be directly activated by temperature changes and certain chemicals applied over their RFs (**Figure 6A**). We did not observe responses to rapid cooling among a sample of lingual LTMRs (n = 8 from 6 mice, **Figure 6B**), consistent with prior findings on the thermal responses of lingual Aδ and Aβ mechanoreceptors in rats (Wang et al., 1993). Moreover, we did not observe responses to chemicals containing the galloyl groups in a series of concentrations, including 100 uM, 200 uM, 500 uM, 1000 uM tannic acid or EGCG applied on the tongue surface (n = 6 from 5 mice, **Figure 6C**). Although hydroxy-alpha-sanshool was shown to be able to interfere with the vibration detection ability of RA channels on the fingertip (Kuroki et al., 2016), we did not find obvious differences in the mean firing rates of lingual RA-LTMRs measured before and after the application of 1300 uM hydroxy-alpha-sanshool (**Figure 6D**). RA-LTMRs also did not respond to the application of a commercial Kava drink on their RFs (n = 6 from 4 mice, **Figure 6E**). This drink contains extracts from the roots of *Piper methysticum* and induces numbing sensations on the human tongue within tens of seconds. Collectively, these results suggest that lingual LTMRs in mice cannot be directly stimulated at the nerve terminals by rapid cooling or certain chemicals that produce astringent or numbing sensations in humans, and are therefore likely unimodal touch sensors.

**Figure 6.**
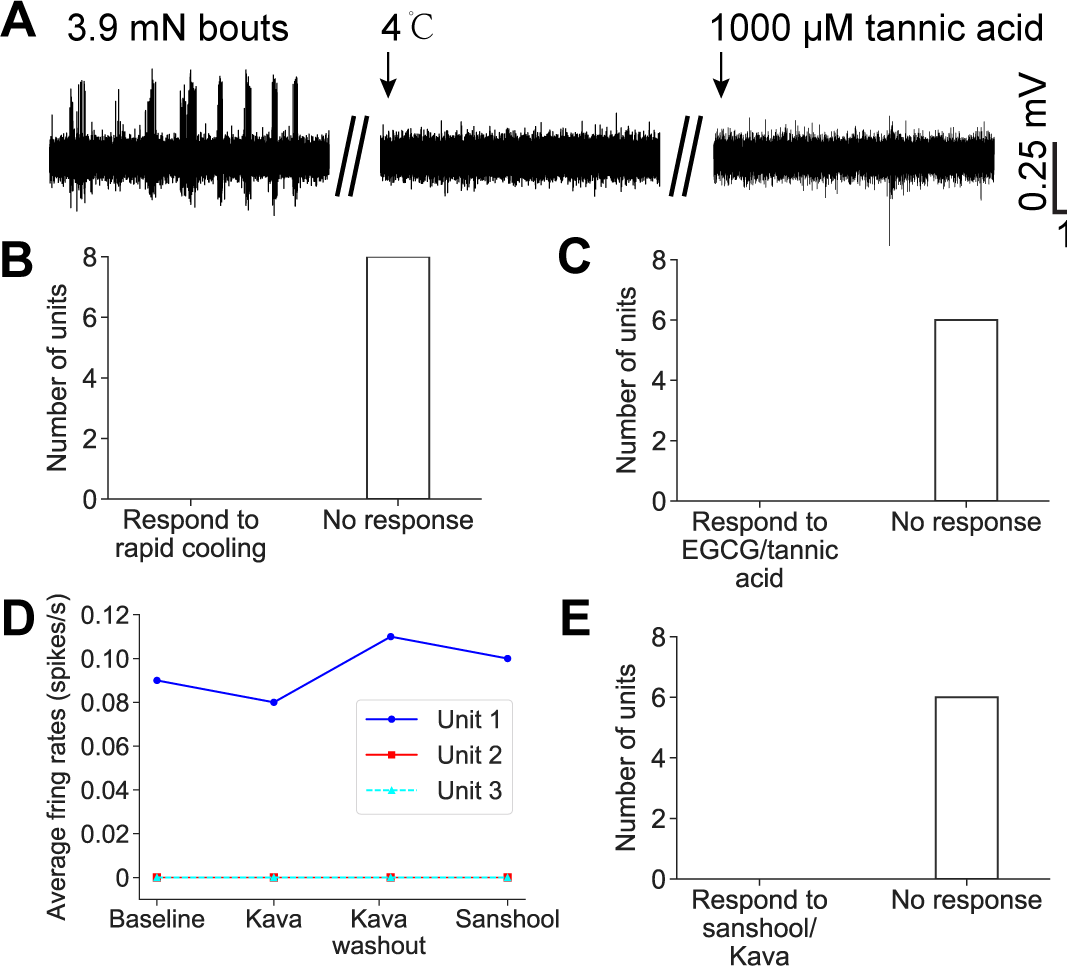
Lingual LTMRs are insensitive to rapid cooling or chemicals that can induce astringent or numbing sensations. **(A)** Neural activities of an example single lingual LTMR unit in TG in response to von Frey indentation, rapid cooling or chemicals that can induce astringent or numbing sensations (only tannic acid at the highest concentration is shown). **(B)** Lingual LTMRs did not respond to rapid cooling (n = 8 from 6 mice). **(C)** Lingual LTMRs did not respond to EGCG/tannic acid in a series of concentrations (n = 6 from 5 mice). **(D)** The average firing rates of 3 example LTMRs before and during Kava/sanshool treatment (from 2 mice). **(E)** Lingual LTMRs did not respond to sanshool or Kava (n = 6 from 4 mice).

## Discussion

In this study, we combined neuronal tracing, genetic labeling and single-unit recording methods to investigate the innervation of the tongue by trigeminal afferents and the response properties of different types of LTMRs in the tongue. We discovered broad innervation of both filiform and fungiform papilla types by TG afferents, and provide evidence that the lingual afferents in the mouse do not contain sensory corpuscles. We found that the adaptation properties of lingual LTMRs fell on a spectrum from classic RA types to IA types, and that tactile sensitivities and preferences differed across the population. Sparse labeling of lingual neurons revealed distinct terminal shapes and branching patterns with regard to the type of papillae they innervate. Together, these results suggest a simple model that relates the physiological properties of lingual LTMRs to anatomically defined patterns of innervation in the dorsal tongue (**Figure 7**).

**Figure 7.**
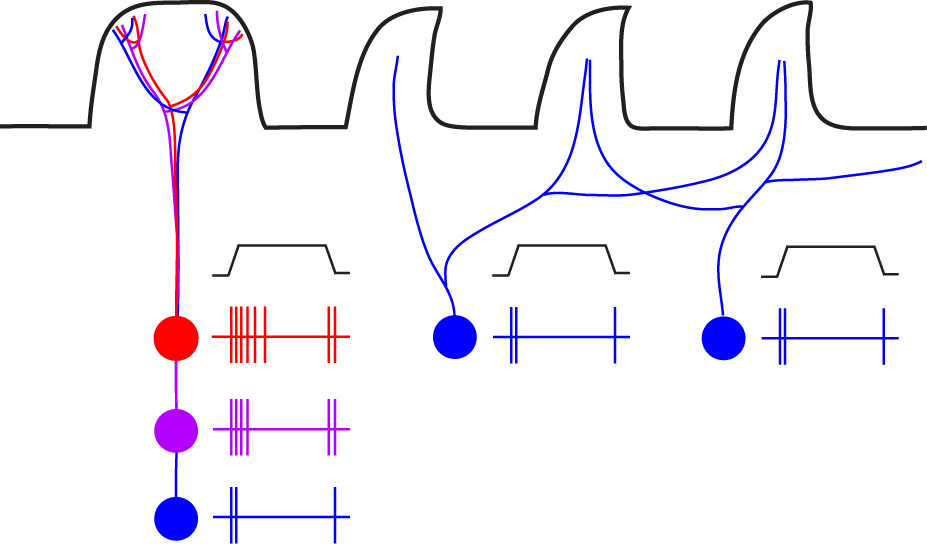
A working model of the organization of tactile innervation in the tongue. The schematic shows a hypothesis of how the lingual papillae at the tongue surface could be innervated by lingual LTMRs in the TG. A single fungiform papilla could be innervated by LTMRs ranging from IA to RA (red, purple and blue), while several adjacent filiform papillae are simutaneously innervated by single RA-LTMRs (blue).

### Fungiform papillae may serve as end organs for touch, providing the tip of the tongue with high tactile sensitivity

The fungiform papillae of the tongue had traditionally been considered as solely gustation-related structures until recently, when mechanosensory afferents in the chorda tympani that respond to stroking and Meissner corpuscle-resembling structures in human tongue were identified (Donnelly et al., 2018; Moayedi et al., 2021). In mice, it has been known that fungiform papillae are innervated by NFH^+^ myelinated afferents and TG afferents (Suemune et al., 1992; Moayedi et al., 2018). However, the relative abundance of TG innervation in fungiform vs filiform papillae remained largely unexplored, as well as the physiological properties of fungiform papillae innervating TG afferents. As a result, it has been assumed for a long time that filiform papillae are the primary mechanosensory organs of the tongue (Lauga et al., 2016).

Our study reveals that compared to filiform papillae, fungiform papillae are more heavily innervated by TG afferents. As TG afferents receiving inputs from fungiform papillae follow a one-to-one projection mode, rather than the one-to-many projection mode for filiform papillae, more neurons are assigned to the innervation of individual fungiform papillae. Roughly, the innervation scheme may be [N > 10^1^ or 10^2^] TG neurons: 1 fungiform papilla, and [N > 10^0^ or 10^1^] TG neurons: [N = 10–20] adjacent filiform papillae. The focalized innervation in fungiform papillae also suggests an emphasis on touch inputs from them in the TG. Indeed, in our physiological study, most lingual LTMRs we recorded from received touch information at the tip of the tongue, where fungiform papillae are most abundant. The locations of most lingual LTMRs’ RFs also resembled those of fungiform papillae, especially those along the midline of the tongue. The small RFs of most lingual LTMRs and the high trigeminal innervation density of fungiform papillae may explain why the tip of the tongue is equipped with high tactile sensitivity. We speculate that fungiform papillae could serve as end organs for touch.

### Comparison between lingual LTMRs and hairy skin LTMRs

The organization of trigeminal nerve endings around two distinct types of papillae on the tongue resembles the way nerves are distributed around hair follicles in the skin. In mice, there are different types of hairs on the skin. Guard hairs, making up ∼1% of hair follicles, are individually innervated by Aβ SA-LTMRs, with each nerve ending terminating into a single guard hair. The more common Zigzag and awl/auchene hairs, accounting for ∼74% and ∼25% of follicles respectively, are innervated by LTMRs whose terminals cover a large area encompassing multiple hairs, including Aβ RA-LTMRs (innervate awl/auchene and guard hairs), Aδ-LTMRs and C-LTMRs (Handler and Ginty, 2021; Kuehn et al., 2019). Similarly, our study revealed that the fungiform papillae, making up ∼2% of lingual papillae, are innervated by potential lingual LTMRs with restricted terminal fields, each of which terminates exclusively at a single fungiform papilla. In contrast, the potential lingual LTMRs innervating the much more abundant filiform papillae (∼98% of papillae) display a more diffused termination pattern, mimicking those of Aβ RA-, Aδ- and C-LTMRs.

Our electrophysiological data suggest that lingual LTMRs have varied adaptation properties, sensitivities and tactile preferences, which might result from the organization of their terminals and the mechanical characteristics of the tissues they innervate. Compared to the more intermediately adapting types, some of the RA types were less sensitive to indentation but equally responsive to brush stroking, resembling RA-LTMRs found in the hairy skin with lanceolate endings. Filiform papillae, being more keratinized than fungiform papillae, likely harbor the least indentation-sensitive lingual LTMRs, which predominantly consist of RA types in the neuronal population. Conversely, LTMRs innervating fungiform papillae are expected to have lower von Frey thresholds (0.08–0.4 mN), including both IA- and RA-LTMRs (**Figure 7**).

In hairy skin, Aβ SA-LTMRs innervating the guard hairs are associated with Merkel cells (Li et al., 2011). In the mouse tongue, however, we did not find any SA-LTMR that could fire throughout the indentation period, except for only a few units with low spontaneous activities. Previous studies have shown that Merkel-neurite complexes may not be a major end organ in the mouse tongue (Moayedi et al., 2018; Donnelly et al., 2022), although some keratin 20-positive cells (potential Merkel cells) associated with NFH^+^ fibers were seen in the base of the ridges between filiform papillae (Donnelly et al., 2022). It has also been known that ablating or dysfunctioning Merkel cells in Merkel-neurite complexes can convert SA afferents into IA afferents (Ikeda et al., 2014; Maksimovic et al., 2014). Therefore, it may not be surprising to see that lingual LTMRs without Merkel cell associations exhibit a spectrum of adaptation properties from RA to IA, but not SA.

### Trigeminal and gustatory touch receptors in the tongue

Previous studies show that GG neurons that can be labeled by the transcription factor Phox2b mainly terminate at the taste bud itself and make contact with taste bud cells, forming intragemmal projections (Ohman-Gault et al., 2017; Tang and Pierchala, 2022). A few Phox2b-positive afferents were also seen to innervate the extragemmal region of fungiform papillae. Those were hypothesized to be mechanosensitive chorda tympani fibers as they were preserved together with the mechanosensitivity of chorda tympani fibers after taste bud elimination via inhibition of the Hedgehog signaling pathway (Donnelly et al., 2022). However, most of the extragemmal afferents in fungiform papillae were not Phox2b-positive, and Phox2b-positive extragemmal afferents were only seen in 43% of the fungiform papillae (Ohman-Gault et al., 2017), suggesting that the extragemmal afferents in fungiform papillae may have a main source other than the GG. We observed that unlike the termination pattern of the GG, the TG afferents terminate at the Piezo2^+^ afferents-enriched extragemmal region of fungiform papillae, which suggests that the majority of the extragemmal mechanosensory afferents have a trigeminal origin. This is consistent with previous findings showing that the chorda tympani denervation had negligible effect on the extragemmal afferents (Nagy et al., 1982; Whitehead et al., 1985).

In summary, TG afferents have a largely different innervation pattern compared to GG afferents. First, TG afferents provide somatosensory innervation to the filiform papillae, which are not supplied by GG afferents. Second, the termination patterns of TG and GG afferents in the fungiform papillae are largely non-overlapping. The TG nerve terminals occupy the Piezo2^+^ afferents-enriched extragemmal space in the apical epithelium, while the majority of GG nerve terminals reside in the intragemmal space, with a few putatively mechanosensitive exceptions. This suggests a functional specialization in the nerve fibers innervating different regions of the fungiform papillae.

### Modality specificity of lingual LTMRs

Previous work has shown that cutaneous C- and Aδ-LTMRs can be stimulated by rapid cooling of the skin (Li et al., 2011). A recent study using *in vivo* calcium imaging of mouse trigeminal neurons also identified lingual mechanosensory neurons that were responsive to a cooling stimulus applied to the tongue surface (Moayedi et al., 2023). Here, our experimental setup and the range of force stimuli we employed were designed in such a way that only lingual LTMRs were recorded, excluding the possibility of recording from potential multimodal nociceptors and neurons innervating deeper tissues. In particular, our sample of lingual LTMRs was insensitive to cooling on the tongue surface, consistent with previous findings showing that both Aδ and Aβ lingual mechanoreceptors in rats are specialized touch sensors with no response to either cold or warm stimuli (8–51 ℃) or menthol (Wang, 1993). This can be explained by the lack of Trpm8 (involved in cooling sensation; Bautista et al., 2007) expression in the vast majority of Piezo2^+^ trigeminal neurons revealed by single-cell RNA sequencing and *in situ* hybridization (von Buchholtz et al., 2021; Nguyen et al., 2017), and lack of mechanical responses in Trpm8-expressing trigeminal neurons (von Buchholtz et al., 2021).

Our sample of tongue LTMRs was not directly stimulated by chemicals that induce astringent or numbing sensations. However, previous work found that some TG neurons can be activated by astringent compounds including tannic acid or EGCG *in vitro* (Schöbel et al., 2014). We think these neurons might be chemosensory or polymodal neurons in TG, and that it is worthwhile to perform *in vitro* chemical tests on Piezo2-expressing TG neurons.

### Limitations of the study and future directions

Although we did not find sensory end organs resembling any known type at the dorsal surface of the mouse tongue, the lingual LTMRs may still be associated with structures that we have not yet examined at their terminals, such as collagen fibers. The potential association of lingual nerve terminals with collagen fibers may also affect their tactile response properties. Therefore, it is worthwhile to study the termination of lingual nerves through careful electron microscopy analysis.

Our model of tongue mechanosensation proposes that LTMRs with low von Frey thresholds (below ∼0.4 mN) innervate the fungiform papillae. This can be directly tested by stimulating a single fungiform papilla during recordings in the TG. In the rat tongue, the fungiform papillae are big enough to enable precise stimulation of a single papilla (Yokota and Bradley, 2016). In mice, although the locations of fungiform papillae can be revealed by staining the tongue with food color dyes, it will require precise apparatus to exclusively stimulate a single fungiform papilla without touching the nearby filiform papillae because of the extremely small sizes of the papillae (**Figure S5**).

In our single-unit recordings, we found a unit with a significantly larger receptive field categorized into the IA type. Our anatomical identification of small lingual afferent terminal sizes could not explain this outlier. It is possible that there are other novel terminal types that we have not found, which occupy a large area, resembling the field LTMRs in the hairy skin (Bai et al., 2015), but exhibit responses to indentation that are different from those of skin field LTMRs.

Finally, our study did not explore the behavioral importance of lingual IA- and RA-LTMRs, and we did not compare the functional importance of TG and GG mechanoreceptors innervating the tongue. In the future, it will be important to modulate the activity of LTMRs by optogenetic activation or inhibition acutely, or to ablate LTMR populations using genetic strategies such as diphtheria toxin receptor (DTR) expression, and to see whether the licking, eating and social-grooming behaviors of the animals are affected. It would also be interesting to study whether a change in the filiform papillae patterning will affect animals’ ability to sense touch information on the tongue by using transgenic mouse lines with altered filiform papillae patterns (Y. Wang et al., 2016).

### Materials and methods Animals

All procedures were in accordance with protocols approved by the Johns Hopkins University Animal Care and Use Committee (protocols: MO18M187 and MO21M195). Mice of both genders were used for experiments. For DiI tracing experiments, mice aged from P0 to P21 from both C57BL/6 and CD1 backgrounds were used. For the other experiments, adult mice aged from 6 weeks to 12 months from C57BL/6 background were used, and the mouse lines used in this study included: *Plp1-EGFP* (*B6;CBA-Tg(Plp1-EGFP)10Wmac/J*), *Vglut2^Cre^* (*B6J.129S6(FVB)-Slc17a6^tm2(cre)Lowl^/MwarJ*), *Rosa26^Ai9^* (*B6.Cg-Gt(ROSA)26Sor^tm9(CAG-tdTomato)Hze^/J*), *Rosa26^Ai32^*(*B6;129S-Gt(ROSA)26Sor^tm32(CAG-COP4*H134R/EYFP)Hze^/J*), *TrkB^CreERT2^* (*B6.129S6(Cg)-Ntrk2/J*), *TrkC^CreERT2^* (*Ntrk3tm3.1(cre/ERT2)Ddg/J*), *Etv1^CreERT2^* (*B6(Cg)-Etv1tm1.1(cre/ERT2)Zjh/J*), *Vglut1^IRES-Cre^* (*B6;129S-Slc17a7tm1.1(cre)Hze/J*), *Advillin^Cre^* (*B6.129P2-Aviltm2(cre)Fawa/J*), *PV^Cre^* (*B6;129P2-Pvalbtm1(cre)Arbr/J*), *TH^Cre^* (*B6.Cg-7630403G23RikTg(Th-cre)1Tmd/J*), *MafA^Cre^* (Xu et al., n.d.), *Pirt^Cre^* (Kim et al., 2008), *Vglut2^Cre^* (*B6J.129S6(FVB)-Slc17a6tm2(cre)Lowl/MwarJ*), *Piezo2-EGFP-IRES-Cre (B6(SJL)-Piezo2tm1.1(cre)Apat/J)* and *Snap25-LSL-2A-EGFP-D (B6.Cg-Snap25tm1.1Hze/J)*. An 8-month-old ferret was used for histological analysis.

To induce CreER-based recombination, tamoxifen was given either in the embryonic stage or to new-born pups. For embryonic stage, A mixture of tamoxifen (3 mg, Toronto Research Chemicals, T006000), progesterone (MilliporeSigma, P0130) and β-estradiol (MilliporeSigma, E8875) in 1:0.5:0.001 ratio prepared in sunflower seed oil (MilliporeSigma, S5007) was administered by oral gavage to pregnant dams. Pups were delivered by Caesarian section at E19-E19.5 and reared by a CD1 foster mother. For new-born pups, 0.1 mg tamoxifen in sunflower seed oil was injected intraperitoneally. The ages of mice when the drug was administered are summarized in **Table 1**.

### Histology

Mice were perfused with PBS transcardially followed by 4% PFA. The tissue was further fixed in 4% PFA at 4℃ overnight, and washed in PBS for 3 times afterwards.

#### H&E staining

The tissue was dehydrated in 30% sucrose in PBS at 4℃ overnight, and protected by OCT embedding compound (4583, Sakura) before frozen. The tissue was then cryo-sectioned into 8-10 µm sections on a Leica cryostat. Sections were hydrated in deionized water, stained in Mayer’s Hematoxylin Solution (MHS1, Sigma-Aldrich) for 15 min, rinsed in warm running tap water for 15 min, then placed in distilled water for 30 s followed by 95% Reagent Alcohol for 30s. The sections were then placed in Eosin Y Solution (HT110216, Sigma-Aldrich) for 30-60 s. Afterwards, they were dehydrated and cleared through 2 changes of 95% Reagent Alcohol, Reagent Alcohol and xylene for 2 min each. The sections were mounted with Balsam Canada Neutral in xylene (846578, Carolina Biological Supply Company), and imaged with a Keyence microscope equipped with brightfield illumination.

#### Masson-Goldner trichrome staining

The tissue was dehydrated in 30% sucrose in PBS at 4℃ overnight, and protected by OCT embedding compound before frozen. The tissue was then cryosectioned into 8-10 µm sections on a Leica cryostat. Sections were hydrated in deionized water, stained with a Masson-Goldner staining kit (1.00485, Sigma-Aldrich). The sections were mounted with Balsam Canada Neutral in Xylene (846578, Carolina Biological Supply Company), and imaged with a Keyence microscope equipped with brightfield illumination.

#### Immunostaining

The tissue was dehydrated in 30% sucrose in PBS at 4℃ overnight, and protected by OCT embedding compound before frozen. The tissue was then cryo-sectioned into 9-40 um sections on a Leica cryostat. Sections were rehydrated in PBS, permeabilized and blocked by 10% goat serum in 0.3% PBST solution (0.3% Triton X-100 (MilliporeSigma, T9284) in PBS) for 1 hour at room temperature, then incubated with primary antibodies diluted in blocking solution (5% goat serum in 0.3% PBST) at 4℃ overnight. The sections were washed by 0.3% PBST for at least 3 times, 15 min each before incubation with secondary antibodies and DAPI diluted in blocking solution at 4℃ overnight. The tissue was finally washed by 0.3% PBST for at least 3 times and PBS for 1 time, mounted with Poly Aqua-Mount (Polysciences, 18606), and imaged using a confocal microscope (Zeiss LSM 880 or Zeiss LSM 700).

#### Whole-mount imaging of DiI-labeled tongue tissue

In P1-P10 mice, after fixation, muscles at the ventral side of the tongue were carefully removed by hand with a surgical blade. The tissue was then mounted dorsal side down on a 30 mm × 10 mm tissue culture dish with 15 mm glass bottom (Celltreat, 229632) in Poly Aqua-Mount. A coverslip was placed on the tissue to gently flatten it. It was then imaged with a wide-field microscope (Zeiss Axio Zoom) or a confocal microscope (Zeiss LSM 880). After whole-mount imaging of the dorsal surface, the tissue was cryo-sectioned. The sections were immediately washed with PBS, mounted with Poly Aqua-Mount and imaged using a confocal microscope (Zeiss LSM 880) after sectioning before DiI diffusion post-cut.

#### Whole-mount imaging of tdTom-expressing tongue tissue post-viral injection

After fixation, muscles at the ventral side of the tongue were carefully removed by hand with a surgical blade. The tissue was then cleared by CUBIC Protocol I as described which can preserve endogenous fluorescent proteins during clearing (Tainaka et al., 2018). Briefly, after fixation and washing of the tissue, the tissue was immersed in CUBIC L solution (10% w/v N-butyldiethanolamine (MilliporeSigma, 471240) and 10% w/v Triton X-100 in water) for 3 days at 37℃. It was then washed in PBS overnight at room temperature, followed by immersion in 50% water-diluted CUBIC R solution (45% w/v antipyrine (Tokyo Chemical Industry, D1876) and 30% w/v nicotinamide (Tokyo Chemical Industry, N0078) in water) at room temperature for 1 day. Finally, it was immersed in CUBIC R solution at room temperature for 1-2 days. All steps were done with gentle shaking. The tissue was imaged with a wide-field microscope (Zeiss Axio Zoom or Olympus BX-41) or a confocal microscope (Zeiss LSM 700). For confocal imaging, the tissue was mounted dorsal side down on a 30 mm × 10 mm tissue culture dish with 15 mm glass bottom in CUBIC R solution. A coverslip was placed on the tissue to gently flatten it.

#### Whole-mount imaging of immunostained NFH

The dorsal surface of the tongue was flattened using a method modified from Y. Wang et al. (2016). Freshly dissected tongue was put dorsal side down into a 3-D printed reservoir and was gently pressed against the bottom of the reservoir by a cell strainer (Fisher Scientific, 22-363-549). 4% PFA was added into the reservoir to fix the tissue at 4 ℃ overnight. The flattened tongue was then fixed for another 1 hour at room temperature, washed in PBS for 3 times, embedded in agarose, and its dorsal part was cut into 300 µm sections using a vibratome (HM 650V, Thermo Scientific). The tissue was then stained and cleared using iDISCO method (Renier et al., 2014). Briefly, the tissue was pretreated with a series of methanol/water (20%, 40%, 60%, 80%, 100%) for 1 hour each, then washed further with 100% methanol (MilliporeSigma, 676780) for 1 hour and chilled at 4 ℃. It was then incubated in 66% dichloromethane (MilliporeSigma, 270997)/33% methanol overnight at room temperature with shaking, washed twice in 100% methanol at room temperature on the next day, and chilled at 4 ℃. Then it was bleached in chilled fresh 5% (v/v) hydrogen peroxide (MilliporeSigma, HX0640) in methanol overnight at 4 ℃, rehydrated with methanol/water series on the next day, and washed in PTx.2 (0.2% Triton X-100 in PBS) for 1 hour at room temperature for 2 times. The tissue was then permeabilized by permeabilization solution (23 g/L glycine (MilliporeSigma, G7126) and 20% DMSO (MilliporeSigma, 472301) in PTx.2) at 37 ℃ for 2 days, blocked by blocking solution (6% normal goat serum and 10% DMSO in PTx.2) at 37 ℃ for 2 days, incubated with 1:1000 Chicken anti-NFH in PTwH (0.2% Tween-20, 10 mg/L heparin in PBS)/5%DMSO/3% normal goat serum at 37 ℃ for 7 days, washed in PTwH at 37 ℃ for 4-5 times until the next day, followed by incubation with 1:500 Goat anti-Chicken IgG Alexa Fluor 647 in PTwH/5%DMSO/3% normal goat serum at 37 ℃ for 7 days and washing in PTwH at 37 ℃ for 4-5 times until the next day. Finally, the tissue was dehydrated with methanol/water series for 1 hour each at room temperature, incubated with 66% dichloromethane/33% methanol for 3 hours with shaking at room temperature, and incubated in 100% dichloromethane for 15 minutes twice with shaking to wash away the methanol. The final dehydration steps and the washing steps were done in a petri dish with the tissue sandwiched by two glass slides to preserve the flattened state of the tissue as much as possible. After that, the tissue was immersed in dibenzyl ether (MilliporeSigma, 108014) at room temperature for refraction index matching until it was cleared, and imaged with a confocal microscope (Zeiss LSM 880).

Primary antibodies used in this study and their dilution ratios or concentration are as follows: Chicken anti-NFH 1:1000 (MilliporeSigma AB5539), Rabbit anti-S100 1:50 (Abcam AB76729), Rabbit anti-S100 1:200 (ProteinTech, 15146-1-AP), Rabbit anti-PGP9.5 1:1000 (MilliporeSigma, AB1761-I), Rabbit anti-NeuN 1:500 (MilliporeSigma, MABN140), Rat anti-Cytokeratin 8 (TROMA-I, DSHB, 2.5 µg/mL).

Secondary antibodies used in this study and their dilution ratios are as follows unless otherwise stated: Goat anti-Rabbit IgG Alexa Fluor 488 1:1000 (ThermoFisher A-11008), Goat anti-Rabbit IgG Alexa Fluor 594 1:1000 (ThermoFisher A-11012), Goat anti-Rabbit IgG Alexa Fluor 647 1:1000 (ThermoFisher A-21244), Goat anti-Chicken IgG Alexa Fluor 488 1:1000 (ThermoFisher A-11309), Goat anti-Chicken IgG Alexa Fluor 647 1:1000 (ThermoFisher A-11039), Goat anti-Rat IgG Alexa Fluor 647 1:1000 (ThermoFisher A-21247).

### DiI tracing

For anterograde DiI tracing from TG to tongue, the DiI crystal labeling procedure was modified from Fei et al. (2014). Briefly, after perfusing mice aged P0 - P21 with PBS and 4% PFA, the brain was removed to expose the TG. The dura covering the mandibular branch of TG was pierced through with #5 forceps. DiI (1,1’-dioctadecyl-3,3,3’,3’-tetramethylindocarbocyanine perchlorate (DiIC_18_(3)), Invitrogen D282) crystals were implanted into the mandibular region on both left and right sides of the TG, and a few drops of 100% ethanol were added onto the region to dissolve DiI. The head, with the tongue and the TG left *in situ* were covered by a wet paper towel for half an hour, then placed in 4% PFA and incubated at 37℃ for 6 months before the tongue tissue was harvested. In another set of experiments (**Figure 2E**), mice aged P10 - P21 were used and their tissue was incubated with DiI at 25 - 37℃ for 18 months.

For retrograde DiI tracing from the peripheral to TG, DiIC_18_(3) crystals were dissolved in DMSO to create a 1 mg/mL solution. This solution was then diluted in PBS at a ratio of 1:5 to achieve a concentration of 0.2 mg/mL. 2.0 µL of the 0.2 mg/mL DiI solution was injected either into the tip of the tongue or the left whisker pad using a 1 µL micro-syringe (Hamilton) for somatotopic mapping. The animals were sacrificed 5-7 days after DiI injection to maximize neuronal labeling. To estimate the distance between the tip of the tongue and the TG, 2.0 µL of the 0.2 mg/mL DiI solution was injected into the tip of the tongue, and the animals were sacrificed at 1 day, 1.5 days, 42 hours, 45 hours, 2 days, and 5 days after the injection.

### Surgery

Adult mice (older than 8 weeks) were implanted with titanium head caps ahead of electrophysiological recordings or viral injections in TG. The anesthesia was induced with 3% isoflurane in O_2_ and then maintained with 1.5% isoflurane. Mice were kept on a heat blanket (Harvard apparatus), and ophthalmic ointment was applied on eyes to keep them moisturized. 0.5% lidocaine was locally injected under the scalp, and dexamethasone (2 mg/kg) was intraperitoneally injected to prevent inflammation. The skin and the periosteum over the dorsal surface of the skull were removed. A headcap was glued by cyanoacrylic gels (Krazy glue) onto the skull and fixed in place by dental cement. Mice were then injected with buprenorphine (1 mg/kg) i.p. for analgesia and allowed to recover for 7 days. Following headcap implantation, a 2.3 x 2.3 mm cranial window was made with the center located at 2 mm posterior to Bregma, 2 mm lateral to Bregma. The exposed brain region was covered with a biocompatible silicone elastomer (e.g. Kwik-Cast, World Precision Instruments) after craniotomy.

### *In vivo* TG electrophysiology

Mice with head caps were anesthetized by intraperitoneal injection of ketamine/xylazine (87.5 mg/kg and 12.5 mg/kg, respectively), head-fixed, and kept on a heating blanket to maintain its body temperature. The tongue was gently pulled out of the mouth with blunt forceps and put on a custom-made stable platform with an angle of 30° from the vertical. The accessible length of the tongue on the platform was 2.5 - 3 mm. The images of the tongue stained by food color dyes were captured with a circuit board USB digital microscope (Chipquick, SMDPMUSB413). To keep the surface of the tongue moisturized, artificial saliva (4 mM NaCl, 10 mM KCl, 6 mM KHCO_3_, 6 mM NaHCO_3_, 0.5 mM CaCl_2_, 0.5 mM MgCl_2_, 0.24 mM K_2_HPO_4_, 0.24 mM KH_2_PO_4_, pH 7.5. Zocchi et al., 2017) at room temperature was applied on the tongue every now and then. A 2 MΩ tungsten electrode (World Precision Instruments, TM33A20) driven by a Sutter manipulator was inserted into the depth of TG (around 5 - 6 mm below brain surface). A reference 2 MΩ electrode was placed outside the craniotomy covered in sterile saline. The differential signal between the recording and the reference electrodes was amplified 10,000x, band-pass filtered between 300 Hz to 3,000 Hz, and acquired at 20 kHz. In order to find tongue units, multiple TG locations surrounding 2.3 mm posterior to Bregma, 2 mm lateral to Bregma were probed. While the electrode was moving down at the depth of TG, the orofacial regions of the mouse including whisker pad, cheek, lower jaw and tongue were touched by a hand-held fine-tip brush in a stroking manner in order to reveal the receptive fields of the units encountered. Once a unit that could respond only to tactile stimuli on the tongue was discovered, it was further tested with von Frey filaments, chemicals, and ice-cold artificial saliva. Von Frey filaments were pointed perpendicular to the RF, and forces ranging from 0.08 mN to 9.8 mN were applied whenever suitable. For units with RFs located away from the edge of the tongue, a range of von Frey stimuli ranging from 0.08 mN to 9.8 mN could be applied. For those with their RFs located at the edge of the tongue, where the local curvature of the tongue surface was significant, their responses to 9.8 mN (and sometimes 5.9 mN) were not tested due to a significant push of the tongue by the larger von Frey filaments. To test the conduction velocity of the unit, a 0.4 mA, 1 ms electric pulse generated by a stimulus isolator (World Precision Instruments, A365) was delivered onto the unit’s receptive field by a concentric bipolar electrode (World Precision Instruments, TM33CCINS). For opto-tagging experiments, 10 ms-long light pulses were generated by a 473 nm DPSS laser (Laserglow Technologies, R471003GX) and delivered to the tongue via a 200 mm, 0.39 NA optical fiber manually positioned around 1 mm away from the unit’s receptive field, with the actual power to be 15 -18 mW. The TTL inputs sent to the stimulus isolator and the laser were generated from a National Instruments DAQ system (BNC-2090A), and were recorded by Wavesurfer (https://wavesurfer.janelia.org/) together with electrophysiological data. When a unit was found with a RF located near the edge of the tongue, it was further probed with tactile and optical stimulus at both ventral and dorsal sides of the tongue to determine its exact RF location. The mice were redosed with 25-50% of the initial ketamine/xylazine once they started to wake up from anesthesia indicated by whisker twitching. The duration of anesthetic state of the mouse between each redose was closely monitored in order to avoid applying excessive doses.

Chemicals applied on the tongue surface include: tannic acid (100, 200, 500, 1000 µM, diluted with artificial saliva, MilliporeSigma, 403040), (−)-epigallocatechin gallate (EGCG, 100, 200, 500, 1000 µM, diluted with artificial saliva, MilliporeSigma, E4143), Hydroxy-α-sanshool (1900 µM, freshly dissolved in DMSO and then diluted with artificial saliva by 1:20. Medchemexpress, HY-N6825), Kava concentrate (commercial drink, Kalm with Kava^TM^, iced tea flavor).

### Viral injection in the TG

Glass pipettes (Wiretrol II, Drummond Scientific Company, 5-000-2010) with a long taper (more than 6 mm) were pulled using a pipette puller (Sutter Instruments, model P-97) for TG injection. The tips of the pipettes were then trimmed by forceps and beveled by gently pressing them against a rotating disk. A hydraulic microinjection system was used for injection. For sparse to dense labeling of mandibular branch neurons, the pipette containing AAVPHP.S-CAG-FLEX-tdTomato virus (Addgene 28306-PHP.S, 1.8×10^13^ vg/mL) was targeted at 1–4 injection sites spaced 150– 200 µm apart in the mandibular branch (lateral 2.3 - 2.5 mm, posterior 2.2 - 2.5 mm relative to Bregma). For each injection site, 0.2 µL virus was injected at a rate of 70 nL/min at a depth between 5.8 to 6 mm. The craniotomy was then covered and protected by silicon elastomers and the mice were sacrificed 3 weeks after viral injection. In another set of experiments, rAAV1-CAG-FLEX-tdTomato virus (Addgene 45282, 1.2×10^13^ GC/mL) was injected into the TG mandibular branch. For dense labeling, a total of 1 - 1.2 µL of virus was injected at 4 locations spaced 150 - 200 µm apart. For sparse labeling, a total of 0.4 - 0.8 µL of virus was injected at 2 locations spaced 200 µm apart. The injection depths were 5.7 - 5.9 mm. The mice were sacrificed 4 - 4.5 weeks after viral injection.

### Piezo2 expression of tongue-innervating neurons

The tongue tissue was collected from *Piezo2-EGFP-IRES-Cre/+;Snap25^LSL-EGFP/+^* mice at P1-P2 after decapitation and was freshly embedded in low-melt agarose (2% at 40 °C) with the dorsal side facing up, followed by imaging with a confocal microscope. To avoid lineage problems caused by embryonic expression of Piezo2, other mice that were *Piezo2-EGFP-IRES-Cre/+;Snap25^LSL-EGFP/+^* or *Piezo2-EGFP-IRES-Cre/+;Snap25^+/+^* were injected with AAV9-pCAG-FLEX-tdTomato-WPRE interperitoneally (10 µL, 6.4×10^12^ GC/mL). The tissue was harvested 14 days after injections. The green autofluorescence and the tdTomato expression at the dorsal surface of the tongue were imaged with a confocal microscope. The autofluorescence signals from the filiform papillae were subtracted from the red channel with Fiji.

AAV9-pCAG-FLEX-tdTomato-WPRE was a gift from Hongkui Zeng (Addgene plasmid # 51503) and was produced by Vigene Biosciences.

### Statistical analysis and data processing

All analyses were performed using custom-written Python or MATLAB (MathWorks) scripts. No statistical methods were used to predetermine sample sizes. The confocal and wide-field fluorescence images were processed using Fiji (ImageJ).

#### DiI intensity across the tongue

The fluorescence intensity of a 500 µm-wide square region-of-interest (ROI) with one edge located 150 - 200 µm away from the tip of the tongue to ensure that the ROI was entirely inside the tongue, and another edge aligned with the location of the farthest fungiform papilla at the posterior of the tongue was analyzed using Fiji “Analyze - Plot Profile”. The averaged fluorescence intensity at each distance along the A-P axis was normalized according to the equation:

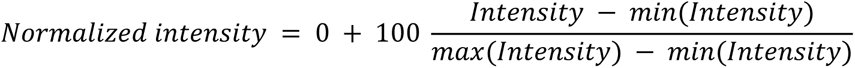

The distance along the A-P axis was also normalized according to the equation:

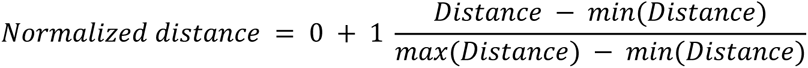

*Innervation intensity surrounding fungiform papillae* Confocal stacks of individual fungiform papillae innervated by DiI-labeled trigeminal afferents were analyzed using Fiji. Both sphere-shaped and ellipse-shaped fungiform papillae were included for the analysis. Image slices containing nerve fibers at the base of fungiform papillae were excluded for maximum intensity projection of the z-stacks. A circle or ellipse was drawn to fit the contour of the papillae, and the image was unwrapped and converted to polar coordinates using the plugin “Polar Transformer” with 360 degrees used for polar space and 1 line per angle. The fluorescence intensity of the outer ring region of fungiform papillae was selected as ROI and analyzed using “Plot Profile”. The fluorescence intensity at each angle for a specific fungiform papilla was normalized according to the equation:

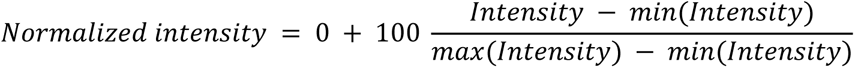

#### Spike sorting

Voltage recordings were high-pass filtered at 300 Hz with custom designed filters using MATLAB R2023a (MathWorks), thresholded, and clustered using MClust 4.4 (made by A. D. Redish, https://github.com/adredish/MClust-Spike-Sorting-Toolbox) to extract single unit waveforms and spike times. Units whose RFs were located at the ventral side of the tongue were disregarded for analysis.

#### Peristimulus time histogram (PSTH) and Adaptation Index

As the exact timing of von Frey hairs’ contact with the tongue could not be registered without changing the force properties of von Frey hairs, the stimulus onset was determined by the timing of the first spike of a unit in response to von Frey stimulation. The 50 ms bin size was adopted after comparing the PSTH results using 10, 20, 30, 50 and 100 ms time bins. The adaptation index (AI_t_) was first calculated using different criteria based on the estimated time taken for the 3.9 mN von Frey stimulus to reach a plateau (t = 150, 200, 250, and 300 ms), after which t = 250 and 300 were adopted. The AI_t_ was calculated by the equation:

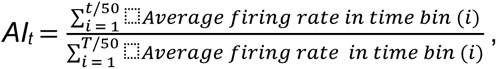

where T was the total duration of a spike train excluding the touch offset responses. Units with a low spontaneous activity (n = 2) were excluded for this analysis.

#### Neuroanatomical tracing and terminal field quantifications

Semi-automatic tracing of sparsely labeled trigeminal afferents was performed using the SNT toolbox (https://imagej.net/plugins/snt/). The terminal field sizes for afferents innervating fungiform papillae were determined by a convex hull enclosing all the endings and the afferent branching point near the fungiform papillae. For afferents innervating multiple filiform papillae, the terminal field included only a narrow area surrounding all the afferent branches near the terminal (not a convex hull).

## Acknowledgements

We thank Xinzhong Dong for providing *Pirt^Cre^* and *MafA^Cre^* mice, Wenqin Luo for providing *Split^Cre^* mice, Kristina Nielsen and Jennifer Smith for providing fixed ferret tongue tissue. We thank Varun Chokshi and Jeong Jun Kim for comments on the manuscript. This work was supported by NIH grants R01NS089652 and 1R01NS104834-01 (D.H.O.) and by the intramural program of the NIH, the National Center for Complementary and Integrative Health, and National Institute of Neurological Disorders and Stroke (A.T.C.).

## Author contributions

L.Z., W.P.O. and D.H.O. planned the project and designed experiments. L.Z. collected and analyzed data under the supervision of D.H.O. M.N. collected the data showing Piezo2 expression in the tongue under the supervision of A.T.C. L.Z. wrote the initial draft of the paper with inputs from all authors. All authors discussed and edited the paper.

## Declaration of interests

The authors declare no competing interests.

---

**Figure S1.**
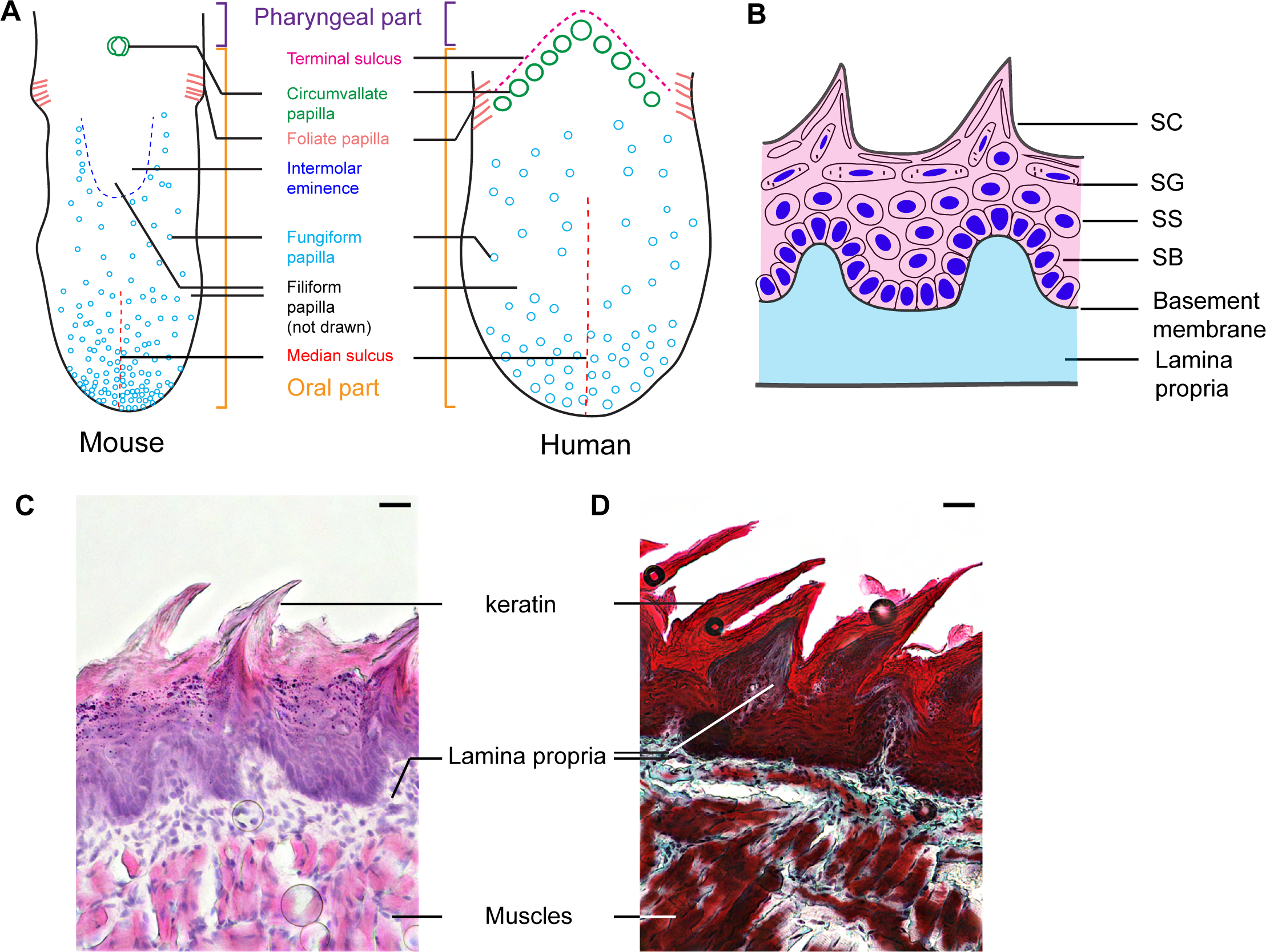
The anatomy of the tongue. **(A)** Left: a schematic outlined from an actual mouse tongue with DiI anterograde tracing from the TG to reveal locations of lingual papillae. Filiform papillae are the most abundant type among the four types of papillae and are distributed across the entire oral part of the tongue, including the intermolar eminence. They are not seen at the pharyngeal part of the tongue posterior to the circumvallate papilla. Fungiform papillae are distributed across the anterior part of the tongue and regions surrounding the intermolar eminence, but not inside or posterior to the intermolar eminence. Right: a schematic of the human tongue modified from Jung et al. (2004). The exact numbers and locations of the lingual papillae (especially the fungiform papillae) may not match those of the real tongue. The sizes of the tongues are out of scale. **(B)** A schematic diagram of the keratinized oral mucosa. SC: stratum corneum. SG: stratum granulosum. SS: stratum spinosum. SB: stratum basale. Blue: cell nuclei. Pink: epithelium. Cyan: lamina propria. **(C)** Lingual mucosa of a 6-week-old mouse stained with H&E. Cell nuclei were stained purple, and proteins were stained pink. Keratin composed of dead epithelial cells was almost transparent. Scale bar: 25 µm. **(D)** Lingual mucosa of a 6-week-old mouse stained with Masson-Goldner staining. Keratin was stained bright red. The lamina propria (connective tissue) stained cyan infiltrated each filiform papilla until reaching the keratin layer. The muscles were stained dark red. Scale bar: 25 µm.

**Figure S2.**
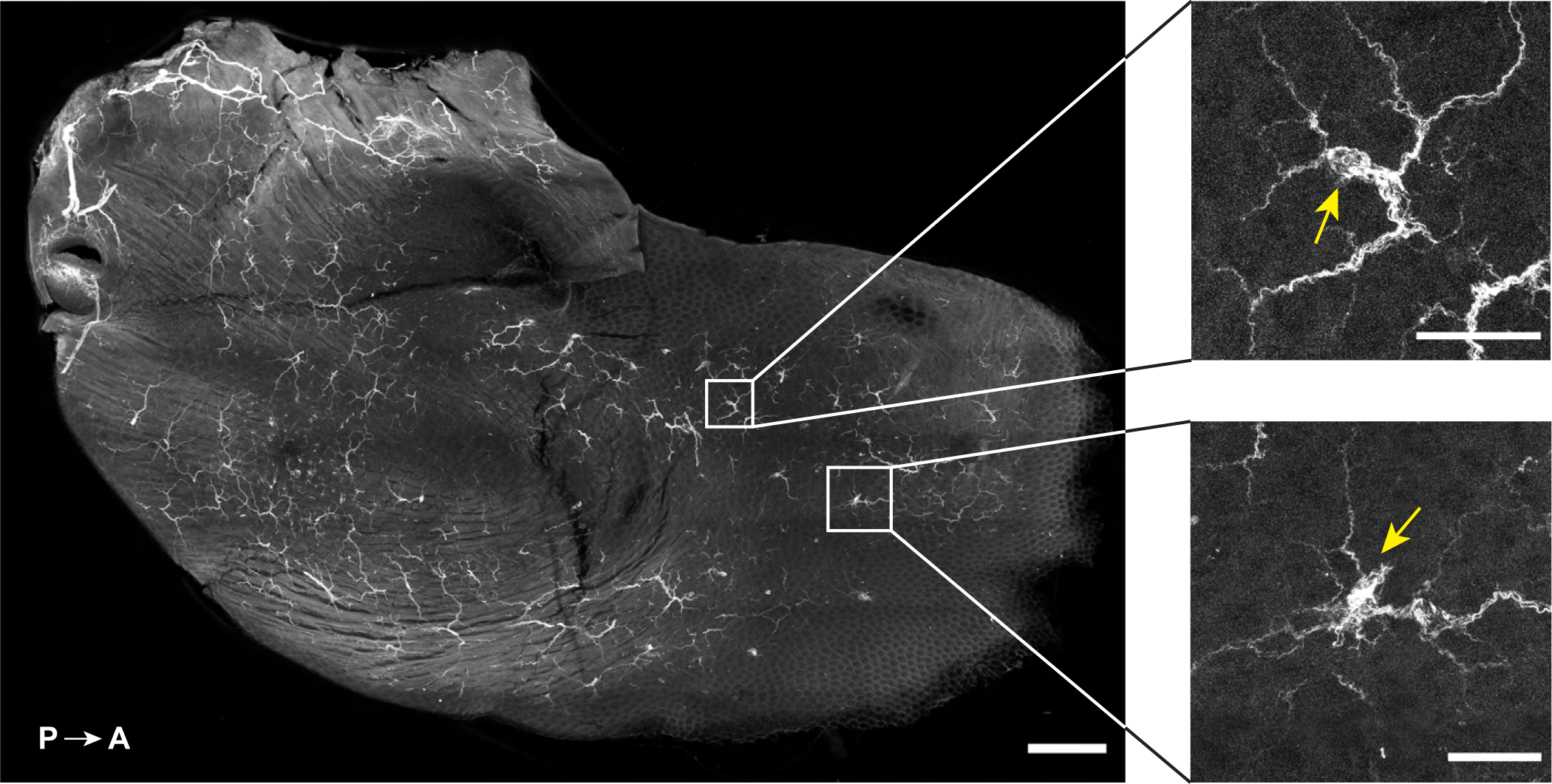
NFH^+^ neural afferents at the dorsal surface of the tongue. Left: dorsal surface of the tongue treated by iDISCO tissue clearing and anti-NFH immunostaining. The tip of the tongue was cut off by sectioning and points towards the right side of the image. NFH^+^ neural afferents innervate both anterior and posterior regions of the tongue dorsal surface, and were seen to be enriched in fungiform papillae (yellow arrows) as shown in insets at the right. The arrows point at fungiform papillae. Scale bars: 500 µm at the left and 100 µm for insets.

**Figure S3.**
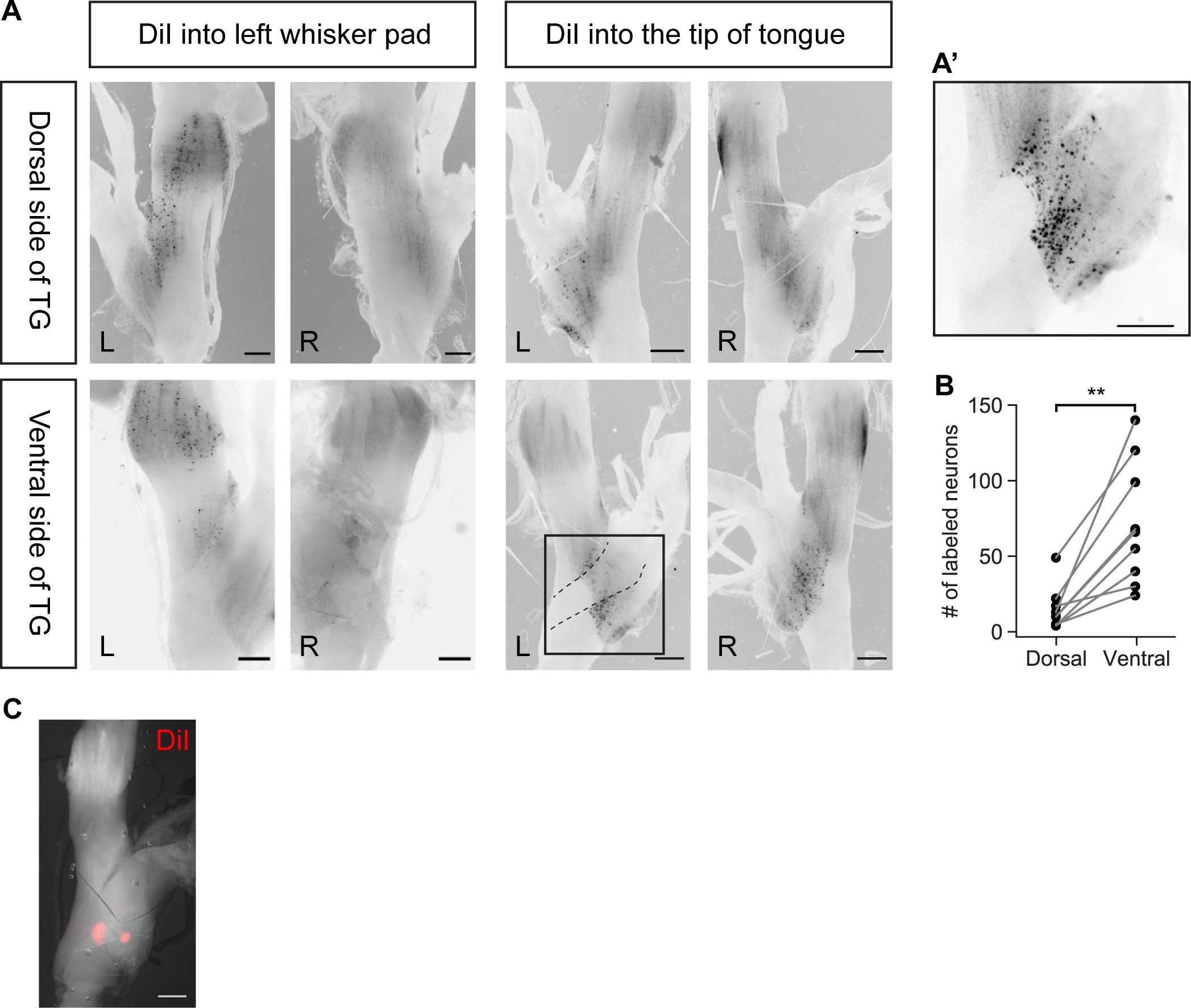
The somatotopic organization of trigeminal ganglion neurons innervating the whisker pad or the tip of the tongue. **(A)** Whole-mount imaging of the dorsal or the ventral surface of TG after DiI tracing from the left whisker pad or the tip of the tongue. The trigeminal neurons innervating the left whisker pad were located in the maxillary branch of the left TG, whereas those innervating the tip of the tongue were in the mandibular branch of TG and were most abundant at the ventral side of TG. The dashed line marks the boundary of a nerve trunk running through the ventral surface of TG. L: the left TG; R: the right TG. Scale bar: 500 µm. **(A’)** A zoomed-in view of the inset in **(A)** after dissecting the nerve trunk to reveal the cell bodies underneath. Scale bar: 500 µm. **(B)** Number of DiI-labeled neurons with a soma size no less than 20 µm that could be visualized with a wide-field fluorescence microscope at the dorsal or ventral side of the TG (n = 9 TG from 5 mice). The nerve trunk at the ventral side was not removed. p = 0.0011, paired t-test. **(C)** The ventral view of the left TG showing location of recorded lingual LTMRs marked by a DiI-coated tungsten electrode. Scale bar: 500 µm.

**Figure S4.**
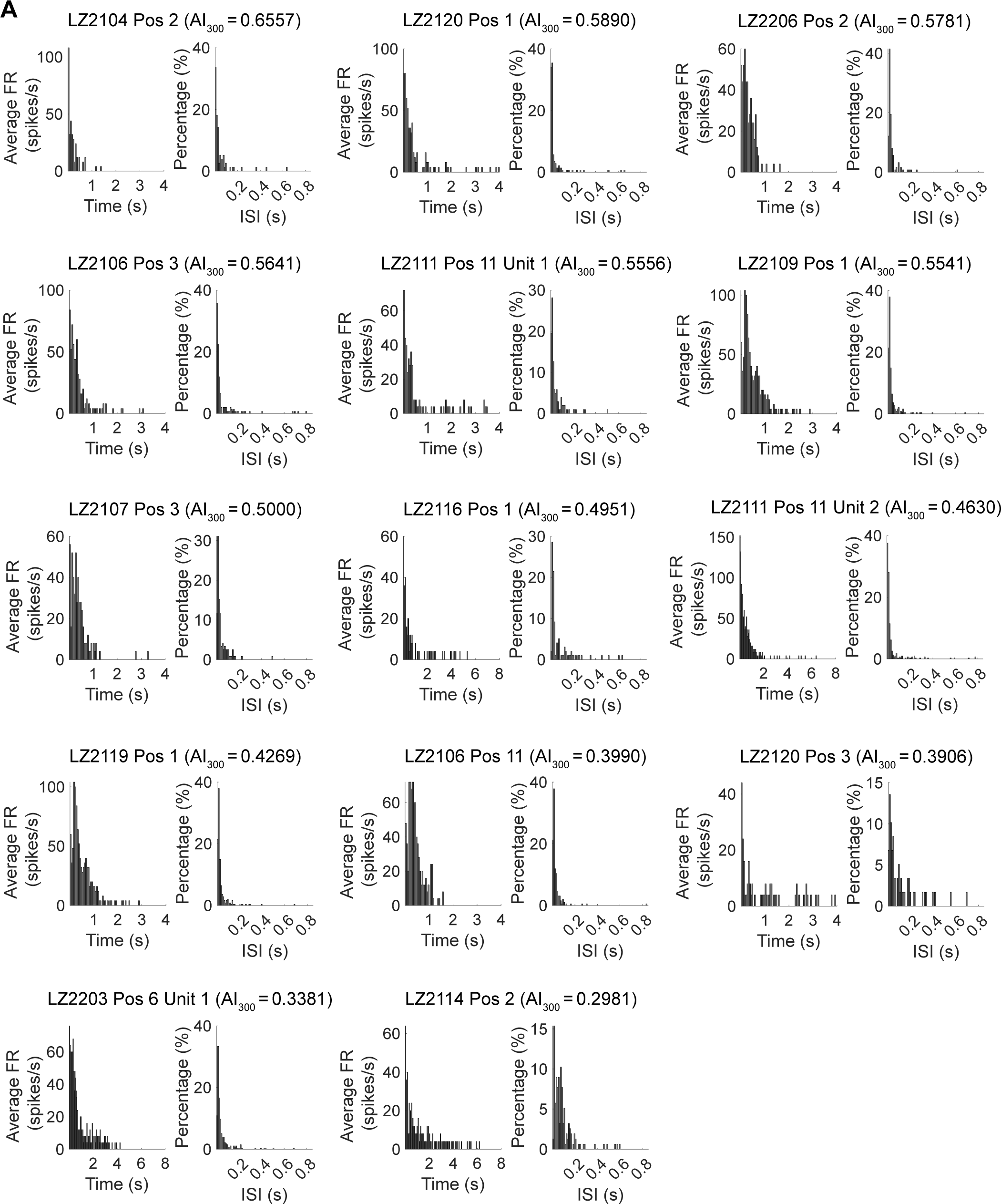
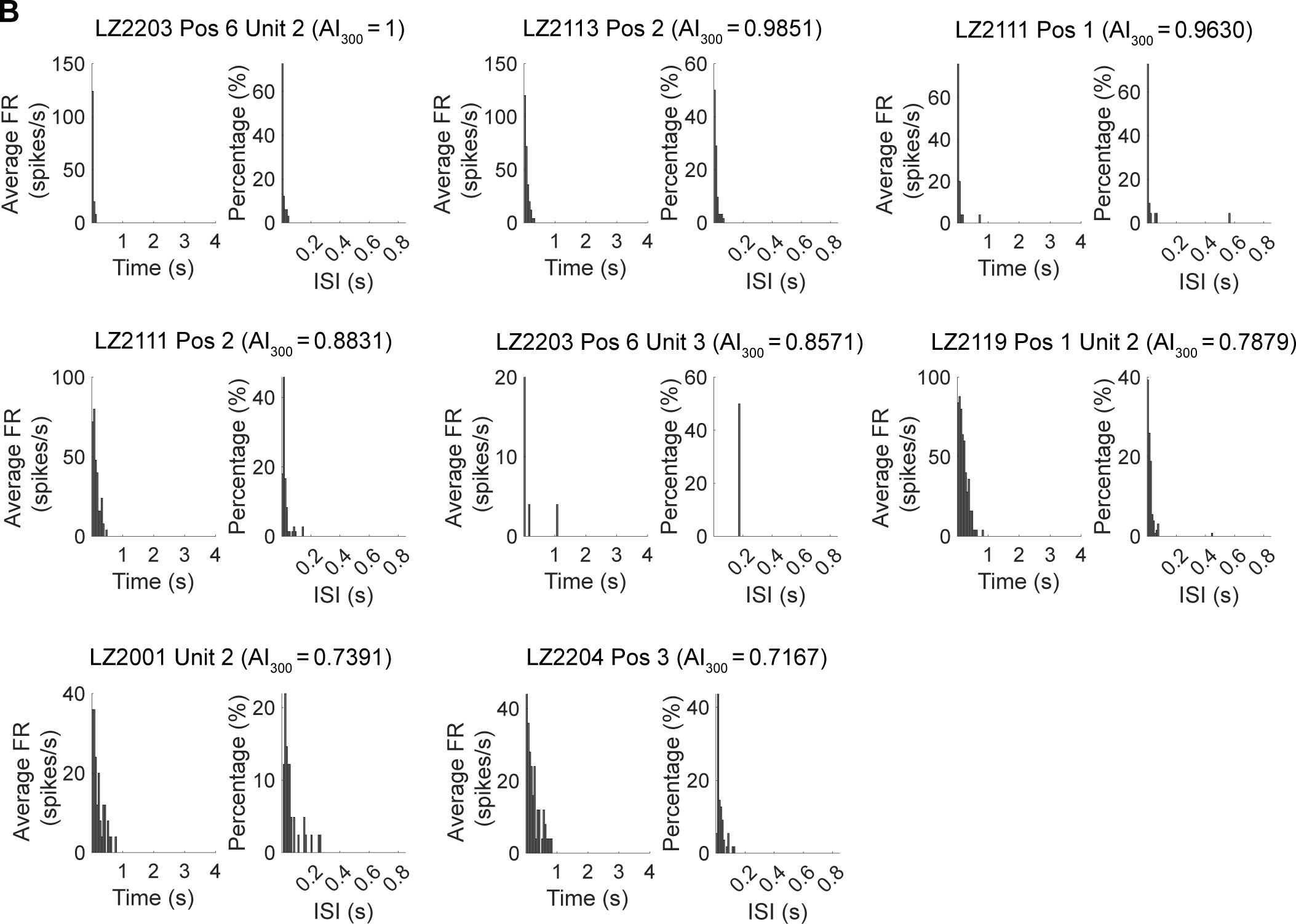
Peristimulus time histograms (PSTHs) and interspike intervals (ISIs) of lingual LTMRs. **(A)** PSTHs and ISIs of IA units sorted with an decreasing AI_300_. **(B)** PSTHs and ISIs of RA units sorted with an decreasing AI_300_. **(A-B)** PSTH bin size: 50 ms, n = 5 events. ISI bin size: 10 ms.

**Figure S5.**
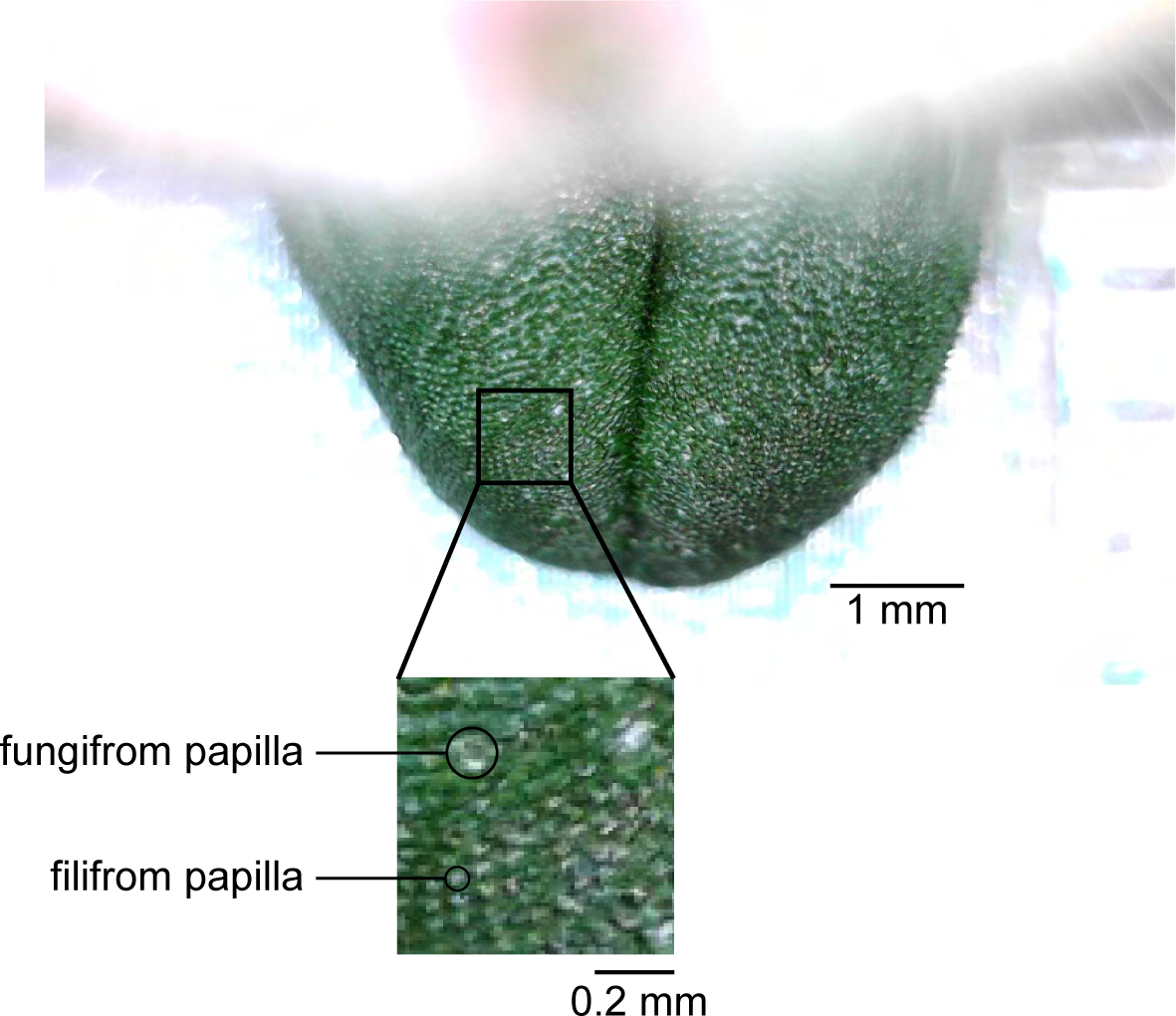
Fungiform and filiform papillae on a mouse tongue stained by a green food color.

## Notes

### Competing Interest Statement

The authors have declared no competing interest.

### Summary of Updates

This version has been revised to include new results from sparse labeling experiments and new analyses of electrophysiological responses.

